# Single-cell profiling of healthy human kidney reveals features of sex-based transcriptional programs and tissue-specific immunity

**DOI:** 10.1101/2021.12.09.471943

**Authors:** Caitriona M. McEvoy, Julia M. Murphy, Lin Zhang, Sergi Clotet-Freixas, Jessica A. Mathews, James An, Mehran Karimzadeh, Delaram Pouyabahar, Shenghui Su, Olga Zaslaver, Hannes Röst, Madhurangi Arambewela, Lewis Y. Liu, Sally Zhang, Keith A. Lawson, Antonio Finelli, Bo Wang, Sonya A. MacParland, Gary D. Bader, Ana Konvalinka, Sarah Q. Crome

**Affiliations:** Toronto General Hospital Research Institute, University Health Network; Toronto, ON, Canada; Ajmera Transplant Centre, University Health Network; Toronto, ON, Canada; Department of Medicine, Division of Nephrology, University Health Network; Toronto, ON, Canada; Department of Immunology, University of Toronto; Toronto, ON, Canada; Department of Statistical Sciences, University of Toronto; Toronto, ON, Canada; Vector Institute; Toronto, ON, Canada; Department of Molecular Genetics, University of Toronto; Toronto, ON, Canada; The Donnelly Centre, University of Toronto; Toronto, ON, Canada; Department of Computer Science, University of Toronto; Toronto, ON, Canada; Department of Laboratory Medicine and Pathobiology, University of Toronto; Toronto, ON, Canada; Peter Munk Cardiac Centre, University Health Network; Toronto, ON, Canada; Princess Margaret Cancer Centre, University Health Network; Toronto, ON, Canada; The Lunenfeld-Tanenbaum Research Institute, Mount Sinai Hospital; Toronto, ON, Canada; Institute of Medical Science, University of Toronto; Toronto, ON, Canada

**Author notes:** co-first authorship (Caitriona M. McEvoy, Julia M. Murphy). equal contribution (Sergi Clotet-Freixas, Jessica A. Mathews). co-corresponding authors: Sarah Q. Crome Ana Konvalinka.

## Abstract

Maintaining organ homeostasis requires complex functional synergy between distinct cell types, a snapshot of which is glimpsed through the simultaneously broad and granular analysis provided by single-cell atlases. Knowledge of the transcriptional programs underpinning the complex and specialized functions of human kidney cell populations at homeostasis is limited by difficulty accessing healthy, fresh tissue. Here, we present a single-cell perspective of healthy human kidney from 19 living donors, with equal contribution from males and females, profiling the transcriptome of 27677 high-quality cells to map healthy kidney at high resolution. Our sex-balanced dataset revealed sex-based differences in gene expression within proximal tubular cells, specifically, increased anti-oxidant metallothionein genes in females and the predominance of aerobic metabolism-related genes in males. Functional differences in metabolism were confirmed between male and female proximal tubular cells, with male cells exhibiting higher oxidative phosphorylation and higher levels of energy precursor metabolites. Within the immune niche, we identified kidney-specific lymphocyte populations with unique transcriptional profiles indicative of kidney-adapted functions and validated findings by flow cytometry. We observed significant heterogeneity in resident myeloid populations and identified an *MRC1^+^ LYVE1^+^ FOLR2^+^ C1QC^+^* population as the predominant myeloid population in healthy kidney. This study provides a detailed cellular map of healthy human kidney, revealing novel insights into the complexity of renal parenchymal cells and kidney-resident immune populations.

## Introduction

The complex functions of the kidney that maintain body homeostasis are executed by a diverse range of specialized parenchymal cells residing in distinct compartments. Within tissues, resident immune populations function in surveillance, maintenance of self-tolerance, response to infection and injury, and interface with parenchymal cells to maintain tissue homeostasis^1–3^. There is limited understanding of this network of kidney parenchymal and resident immune cells in humans due to lack of access to healthy, fresh tissue. Much of our knowledge is based on studies that used kidneys rejected for transplant or tumour-adjacent nephrectomy specimens, where parenchymal populations can have altered molecular programs, and immune populations and their signalling circuits may not be entirely reflective of the steady-state^4, 5^. Further, sex-based dichotomy in gene expression within human kidney cell populations has not been thoroughly examined, but is of great significance to acute and chronic kidney disease, ischemia-reperfusion injury (IRI) and progression of diabetic kidney disease, which exhibit a male preponderance^6–8^.

Here we present a detailed atlas of healthy human kidney using single cell RNA sequencing (scRNAseq) of living donor kidney biopsies, capturing parenchymal and immune cell transcriptomes reflective of a healthy state. We explore sex-based dichotomy in gene expression among kidney populations, revealing altered transcriptional programs between male and female proximal tubular cells, and perform an in-depth characterization of the immune niche in healthy, non-inflamed kidney.

### Single-cell map of healthy human kidney

We examined the cellular landscape of human kidney using pre-implantation kidney biopsies from 19 sex-matched living kidney donors (**Fig. 1a, b**). Our dissociation method was developed to maximize viability to preserve representation of rare and fragile cell populations, and we employed rigorous quality control. Minimal immune cell representation in healthy kidney (∼0.3% of cells captured) necessitated CD45-enrichment for immune cells in 10/19 biopsy samples (5 female, 5 male) (**Fig. 1a).** Of 27677 cells in our map, 6899 cells were from CD45-enriched samples, while 20778 cells were from non-CD45-enriched samples. Twenty-three clusters were identified including several distinct immune cell populations, alongside all anticipated parenchymal populations of the nephron (**Fig. 1c, Supplementary Fig. 1a**). Clusters were comprised of cells captured from multiple donors, there was no exceptional variability in cell cycle state across clusters, and most clusters had symmetrical distribution of donor sex (**Figure 1b-e**, **Supplemental Figure 1**).

**Figure 1.**
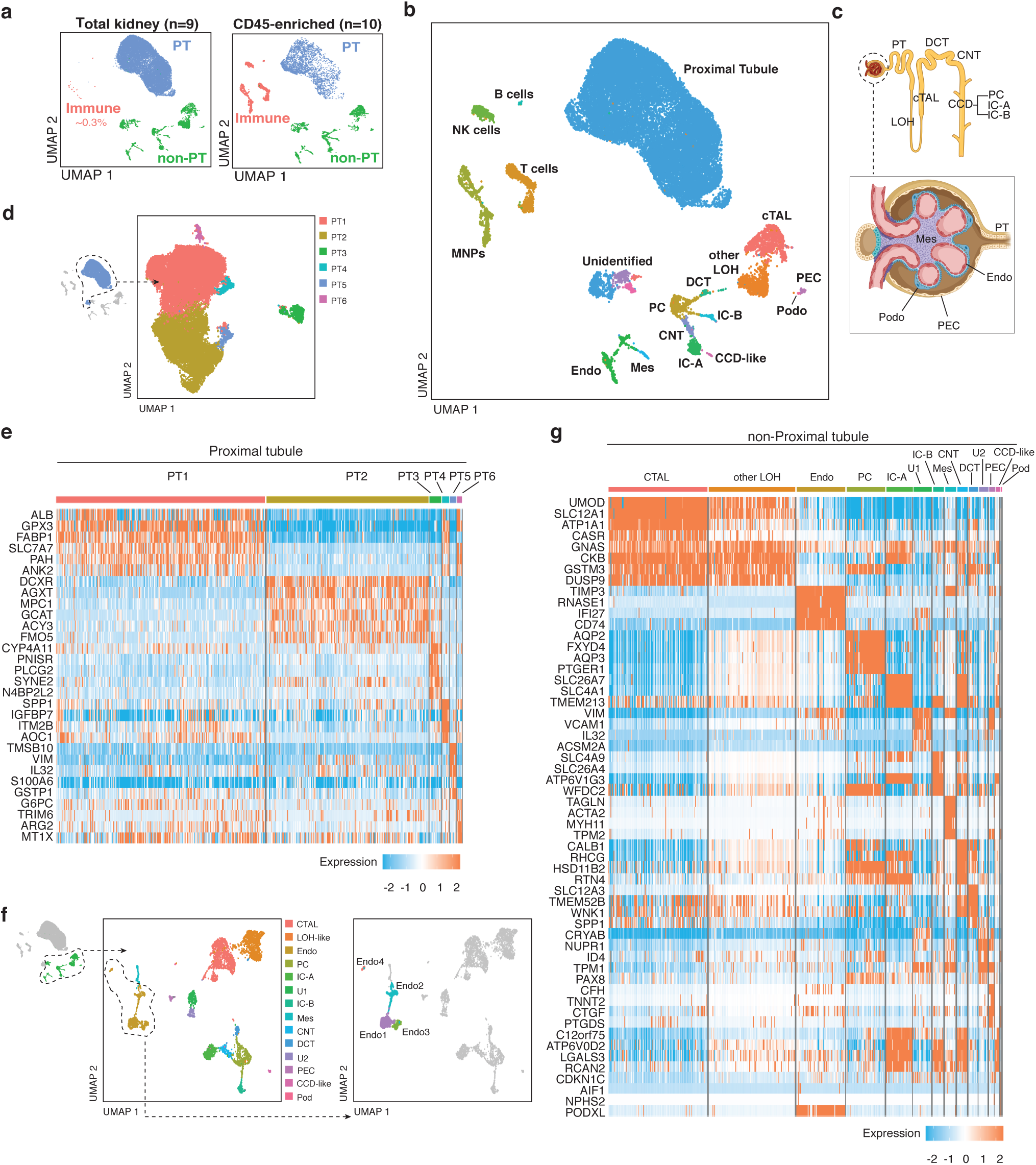
Identification and annotation of kidney parenchymal cells. (**a**) Different cell type proportions were captured by sequencing total kidney homogenate and CD45-enriched samples to create the total combined dataset. (**b**) UMAP clustering of total combined dataset with cell type annotations. (**c**) Graphical depiction of location of nephron cell types captured within the data. (**d**) UMAP plot of compartment-specific analysis of 20772 proximal tubular cells, comprising 6 clusters. (**e**) Heat map showing the expression levels of cluster marker genes. (**f**) UMAP plot of compartment-specific analysis of 4436 non-proximal tubular parenchymal cells, with 14 cell populations represented, including four distinct endothelial clusters. (**g**) Heat map showing the expression levels of cell type marker genes across the 14 non-PT cell populations.

As anticipated, Proximal Tubular (PT) cells comprised 75% of sequenced cells. Sub-clustering revealed 6 distinct clusters (PT1-PT6) (**Fig. 1d, Supplementary Fig. 2a**), with some heterogeneity between individuals, methods of sample preparation, and sexes noted (**Supplementary Fig. 2b**). PT segment-specific separation is evident; PT1, 4, and 6 are enriched for PT segment 1 (S1) marker *SLC5A2* and S1/2-abundant genes (*SLC7A7*, *ANK2*, *SLC4A4*, *SLC6A19*, *SLC22A8*), while PT2 shows increased expression of S3-abundant genes (*DCXR*, *AGXT*, *SLC22A7*, *SLC7A13*) (**Fig. 1e**)^9, 10^. PT3 highly expresses dissociation stress-associated genes^3^, together with general (*LRP2*, *CUBN*) and segment-specific PT genes, indicating cell contributions from all PT segments (**Fig. 1e, Supplementary Fig. 2c**). PT5 (*VIM^+^S100A6^+^VCAM1^+^DCDC2^+^ANXA4^+^)* displays similarity to a putative regenerative PT population – termed ‘scattered tubular cells’ (STC)^11, 12^. These genes also characterize a population which expands following IRI and is postulated to reflect failed PT repair, though expression was also observed in healthy kidney^13^. Some STC-associated genes were exclusively expressed by PT5 or PT3, while others were expressed in both populations (**Supplementary Fig. 2d-f**). This transcriptional overlap between the regenerative STC-like PT5 and stressed PT3 cells may indicate attempted initiation of repair in PT3 cells. Transcription factor analysis (**Supplementary Fig. 2g**) of PT5 genes revealed potential upstream regulators directing cell differentiation and migration (SNAI2, ZNF217), and epithelial phenotype maintenance (ELF3), alongside *NFE2L2* (NRF2), a key regulator of antioxidant and cytoprotective genes^14^. Predicted upstream regulators for PT3 (EGR1, FOS, and JUN) are associated with oxidative stress and fibrogenesis. Predicted regulator ATF3 (protective in renal IRI^15^) supports potential reparative processes in this cluster (**Supplementary Fig. 2g**).

Fourteen non-PT parenchymal cell populations were identified^9^ (**Fig. 1f-g**) including rare but important glomerular populations such as podocytes, mesangial cells, and parietal epithelial cells. We detected notable heterogeneity in CTAL and endothelial populations. Two CTAL subpopulations expressing *CLDN10* and *CLDN16,* respectively, identify cells with differing paracellular cation-resorption preferences in CLDN10-dominant (Na^+^) versus CLDN16-dominant tight junctions (Ca^2+^, Mg^2+^) (**Supplementary Fig. 3a-e**)^16^. Among endothelial subpopulations (Endo1-4) (**Fig. 1f**), we identified two populations (Endo1, Endo3) of peritubular capillary cells (*PLVAP*^+^*TMEM88*^+^*DNASE1L3^+^*) (**Supplementary Fig. 3a, f, g**). Endo1 expressed *ESM1* – required for VEGF-related maintenance of the peritubular capillary network^17^, while Endo3 expressed motility and angiogenesis markers *MARCKS*, *CLU*, *ACKR1*, *SEMA3D* (**Supplementary Fig. 3f**). Endo2 (*SOX17*^+^*SERPINE2*^+^*CLDN5*^+^*CXCL12^+^)* represents afferent arterioles and vasa recta, exhibiting reduced *KDR* expression and increased expression of extracellular matrix-encoding genes (**Supplementary Fig. 3g**). Endo4 expresses the glomerular microvascular endothelial cell markers *EDH3, SOST,* and *TBX3*, a transcriptional regulator critical to fenestrated glomerular endothelial development (**Supplementary Fig. 3f**)^18^.

### Identification of sex-based transcriptomic differences in proximal tubular cells

Leveraging the sex-balanced large sample size, we examined differences in gene expression in healthy human kidney between males and females. Using varimax-rotated principal component analysis, we examined individual kidney populations for separation due to donor sex, and observed a clear separation for the PT population (left panel in **Fig. 2a, Supplementary Fig. 4a**). Such separation was not evident in other cell populations, perhaps reflecting insufficient power with fewer cells. Consequently, subsequent analyses focused on PT cells. Using machine learning, we identified the most discriminant subset of genes in our dataset that could correctly classify cell sex. Model-1 (80 genes) correctly classified cell sex with an area under the curve (AUC) of 0.98 (training dataset), and an accuracy of 84% (validation dataset) (middle panel in **Fig. 2a, Supplementary Fig. 4b,c,f**). As X- and Y-linked genes potentially drive sex-biased effects^19^, we removed all sex chromosome-encoded genes and derived Model-2 (15 genes), which correctly classified cell sex in the training dataset (AUC 0.85), but had reduced accuracy (68%) in the validation set (**Supplementary Fig. 4d-f**). Using an independent single-cell kidney dataset for validation^20^, our gene signatures accurately classified cell sex in 79% (Model-1) and 66% (Model-2) of cells (**Supplementary Fig. 4f**). Next, we identified genes with significant differential expression between males and females (n=75 genes, p-value <0.05, LogFC>0.25) (right panel in **Fig. 2a**). As our conservative analysis excluded genes expressed uniquely by one sex (e.g. Y-chromosome-encoded genes), these genes (n=12) were added for downstream analyses (**Fig. 2b**). Results from our three analyses were compared (**Supplementary Table 1**). In agreement with previous studies^21, 22^, the majority of the sex-biased genes uncovered are located in autosomes, rather than in sex chromosomes. Several sex-biased genes are consistent with previous reports of genes upregulated in murine male (*NAT8, FKBP5, KDM5D, DGKG*) and female (*MGST3, SLC3A1, CYP4A11, RPS29*) PT cells, respectively^21–23^.

**Figure 2.**
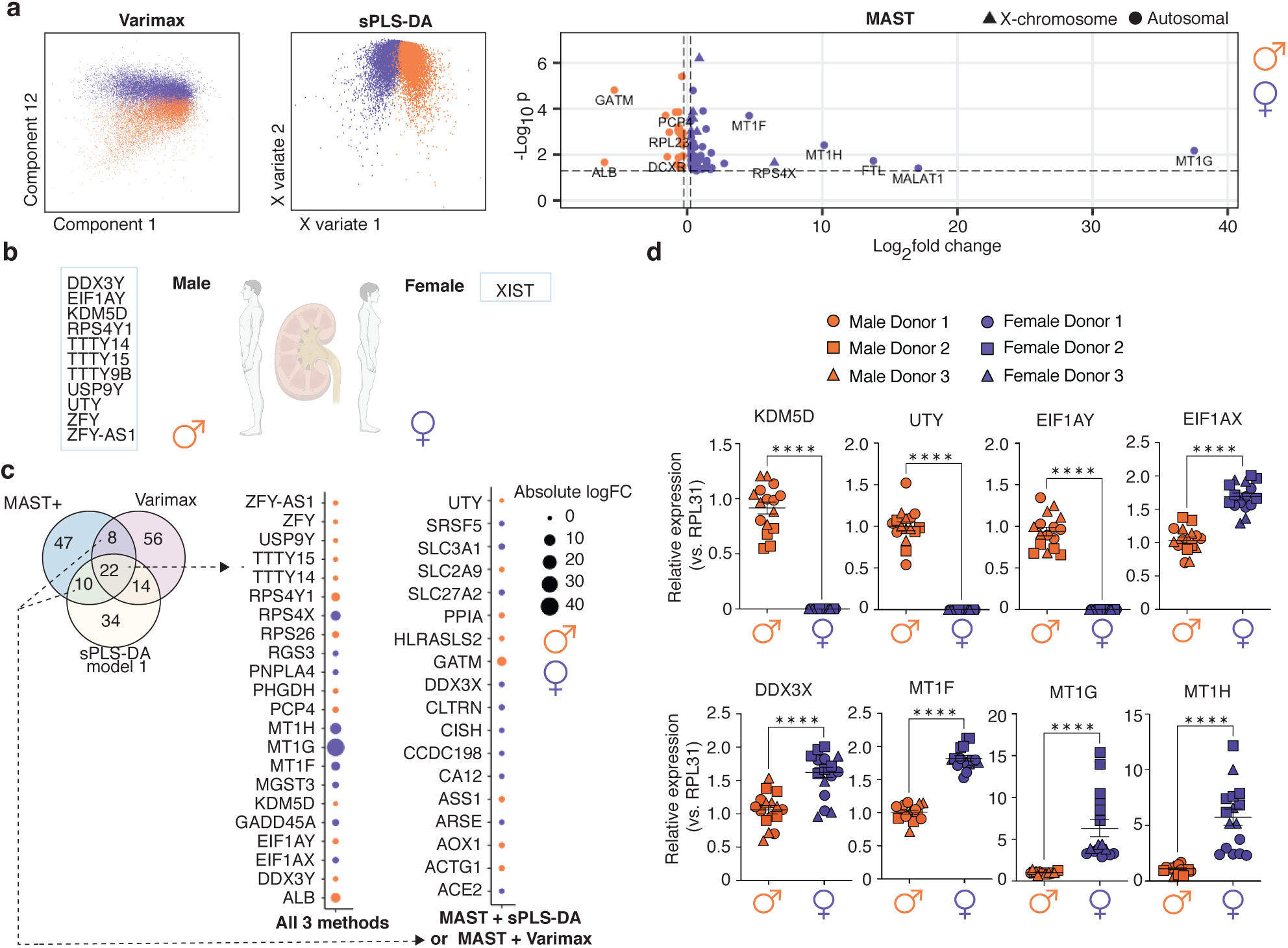
Identifying genes differentially expressed between male and female proximal tubular cells. (**a**) 2-Dimensional plots of Varimax-rotated PCA and sPLS-DA showing separation of male and female cells, and volcano plot showing differential expression of genes between sexes from MAST analysis with sample random effect. (**b**) Genes expressed exclusively by all samples of one sex and none of the opposite sex, which were added to the MAST results for comparison across methods in c. (**c**) Venn diagram depicting genes identified through each analysis, with bubble plots highlighting genes identified by all three methods or by MAST plus one additional method. The size of the circle is proportional to absolute logFC and the colour indicates whether the gene was higher in male (orange) or female (dark purple) PT cells. (**d**) Differences in gene expression of *KDM5D* (p<0.0001, t=17.32, df=30)*, UTY* (p<0.0001, t=18.75, df=30), *EIF1AY* (p<0.0001, t=18.04, df=30), *EIF1AX* (p<0.0001, t=9.077, df=29), *DDX3X* (p<0.0001, t=5.619, df=29), *MT1F* (p<0.0001, t=16.04, df=30), *MT1G* (p<0.0001, u=0), and *MT1H* (p<0.0001, t=6.286, df=30) were determined in primary male and female PT cells, and normalized to RPL31 (n=3 donors/sex; n=4-6 replicates/donor). Group-to-group differences were assessed using two-tailed unpaired t-tests for variables following a normal distribution, and Mann-Whitney tests for variables with a non-parametric distribution. ****p<0.0001. PT, proximal tubule.

Twenty-two genes featured in all three analyses (**Fig. 2c**), including 9 Y- and 3 X-chromosome encoded genes. An additional 18 genes featured in differential expression analysis (MAST+) and one other analysis (**Fig. 2c**). The X-chromosome genes reported are known to escape X-chromosome inactivation, explaining their higher expression in females^19^. Many of the autosomal-encoded genes or their family members are associated with primary sex determination (*SRSF5*^24^, *GATM*^25^, *GADD45A*), sex-biased expression (*CISH*, *SRSF5*, *ACTG1*, *GATM, AOX1*), or sex-specific effects (*SLC2A9*^26^). Intriguingly, many of the genes have established links with kidney disease, including *SLC27A2* (diabetic kidney disease)^27^, *SLC3A1* (cystinuria), and *GATM*^28^; while others are associated with hypoxia (*PHGD, CA12*), inflammation (*PPIA*), and genotoxic stress (*ASS1*). Metallothionein gene family members (*MT1F*, *MT1G*, *MT1H*), which encode cysteine-rich antioxidant proteins^29^, were notably higher in females (**Fig. 2a,c**). Additional differentially expressed genes also relate to cysteine-glutathione availability and metabolism, including *SLC3A1*^30^, *MGST3*, and *HRASLS2*.

We next aimed to validate sex-biased gene expression profiles using commercially-available human primary PT cells from 3 male and 3 female independent healthy donors (**Supplementary Table 2**, age range of donors 50-59 years old). As expected, Y-linked genes *KDM5D*, *UTY*, and *EIF1AY* were exclusively expressed in male PT cells (**Fig. 2d**). We also studied the X-linked genes *EIF1AX* and *DDX3X*. While proposed as ‘X-inactivation escapees’, the extent of X-inactivation can be highly variable across genes, tissues, and individuals^31^. In agreement with our scRNAseq findings, primary female PT cells displayed increased transcript levels of *EIF1AX* and *DDX3X*, compared to male cells (**Fig. 2d**). Female sex is linked to lower oxidative stress markers in the kidney *in vivo*^6^ but whether the sex of PT cells is a major contributor to this effect is unknown. Gene expression of *MT1F, MT1G, MT1H* was significantly increased in primary female PT cells, compared to male cells, as identified by scRNAseq and validated with qPCR in these independent donors (**Fig. 2a,c,d**). Of note, many of the transcripts exhibiting sex dimorphism in our scRNAseq analysis were absent when using matched single nucleus RNA sequencing, likely due to cytosolic or mitochondrial localization of the transcripts (**Supplementary Fig. 5**)

We next investigated the biological processes enriched among the genes showing sex-biased expression in PT cells. Pathway analysis (**Fig. 3a**, **Supplementary Table 3**) revealed processes related to amino acid metabolism, PT transport, and regulation of the inflammatory response as increased in females. Among the pathways increased in males, processes related to mitochondrial aerobic metabolism (‘oxidative phosphorylation’, ‘tricarboxylic acid (TCA) cycle’ and ‘electron transport chain’) predominated. Two additional metabolic processes, namely ‘generation of precursor metabolites’ and ‘nucleoside triphosphate metabolism’, were also enriched in males. To validate these observations, we studied functional differences in mitochondrial metabolism and precursor metabolite generation in male and female PT cells. We exposed primary male and female PT cells to minimal media containing glucose and glutamine, which serve as mitochondrial substrates. We then measured their oxygen consumption rate (OCR), as a marker of mitochondrial respiration^32^. Supporting our pathway analysis, male PT cells showed a significant increase in OCR at baseline and after metabolic stress, compared to female PT cells (**Fig. 3b**). By calculating the corresponding areas under the OCR curves, we determined that male PT cells had a significantly higher basal respiration, ATP-linked respiration, maximal respiratory capacity, and reserve capacity than female cells (**Fig. 3c**). Together with mitochondrial respiration, glycolysis is a major mechanism of glucose-derived energy production^33^. Thus, a parallel increase in glycolysis and aerobic respiration is often indicative of a higher energy state^34^. Increased OCR in our male PT cells was linked to a significant increase in their glycolytic capacity (**Supplementary Fig. 6**), suggesting that they are energetically more active than female PT cells. Mitochondrial respiration results in the generation of two key energy precursors - NAD and ATP^35^. In line with increased aerobic metabolism, male PT cells exhibited a significant increase in the intracellular levels of NAD, β-nicotinamide mononucleotide (NAD precursor), ATP, and three additional nucleoside triphosphate metabolites - GTP, ITP, and UTP (**Fig. 3d**).

**Figure 3.**
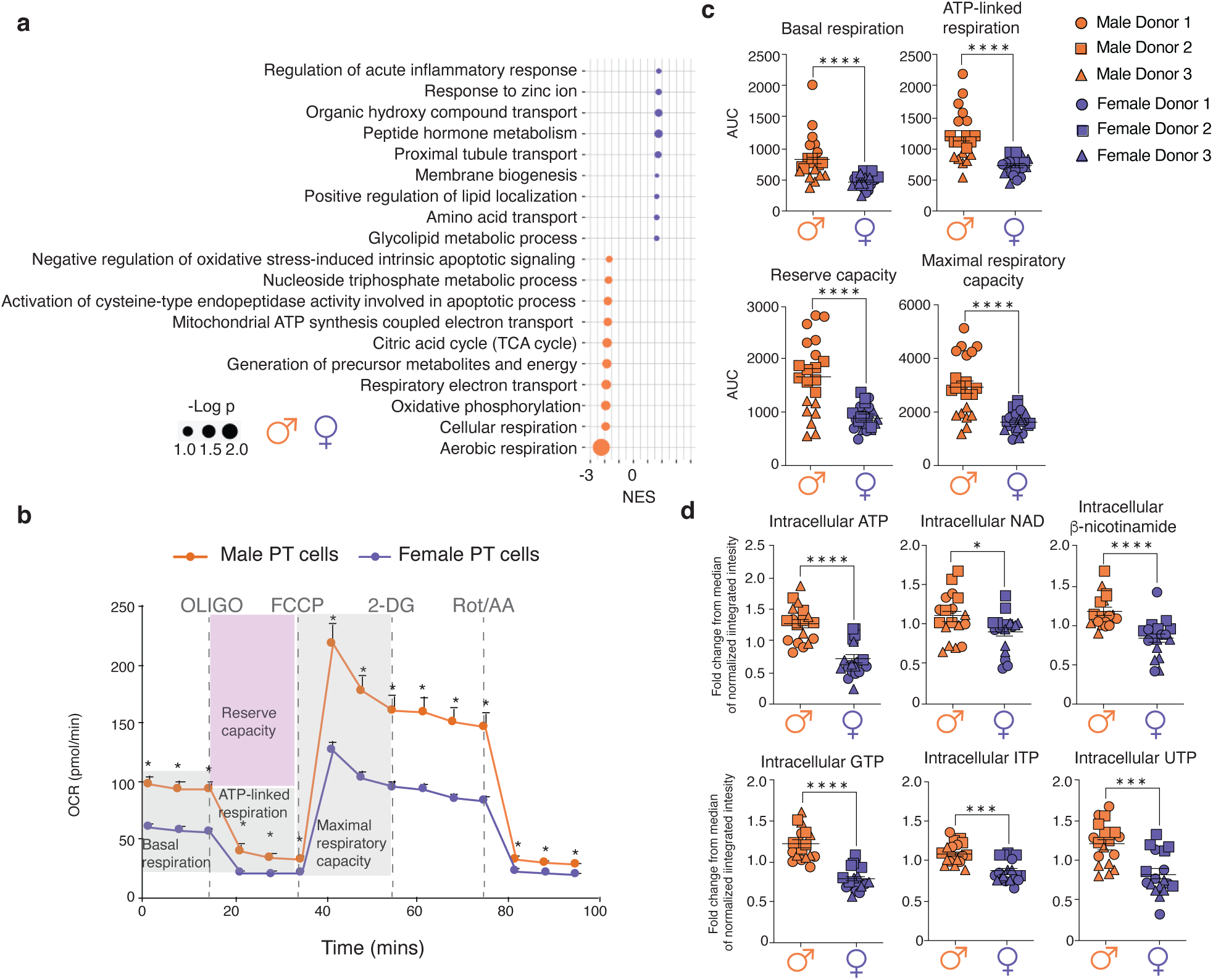
Sex differences in the mitochondrial respiration and energy precursor metabolism of proximal tubular cells. (**a**) Depiction of selected significant (FDR<0.25) terms identified by GSEA analysis as being enriched in males and females respectively. (**b**) Oxygen consumption rate (OCR) was monitored to assess the mitochondrial respiration of male and female PT cells at baseline and after metabolic stress. To induce metabolic stress, the following sequence of drugs was injected: 1μM oligomycin, 0.3μM FCCP, 100mM 2-DG, 1mM Rot/AA. The OCR was monitored in male and female PT cells (n=3 donors/sex; n=6-8 replicates/donor). (**c**) The basal OCR (p<0.0001, u=48), ATP-linked respiration (p<0.0001, t=5.223, df=42), reserve capacity (p<0.0001, t=5.018, df=42) and maximal respiratory capacity (p<0.0001, t=5.281, df=42) of male and female PT cells were calculated from the OCR curves in panel B. Group-to-group differences were assessed using two-tailed unpaired T tests for variables following a normal distribution, and Mann-Whitney tests for variables with a non-parametric distribution. (**d**) In a separate experiment, the intracellular levels of ATP (p<0.0001, t=5.959, df=34), NAD (p=0.029, u=93), β-nicotinamide mononucleotide (p<0.0001, t=4.575, df=34), GTP (p<0.0001, t=7.45, df=34), ITP (p=0.0001, u=46), and UTP (p=0.0001, t=4.316, df=34) were determined in male and female PT cells (n=3 donors/sex; n=6 replicates/donor). Group-to-group differences were assessed using two-tailed unpaired T tests for variables following a normal distribution, and Mann-Whitney tests for variables with a non-parametric distribution., *p<0.05;**p<0.01;***p<0.001;****p<0.0001. PT, proximal tubule; AUC, area under the curve; OCR, oxygen consumption rate; FCCP, p-trifluoromethoxy carbonyl cyanide phenyl hydrazone; 2-DG, 2-deoxyglucose; Rot, rotenone; AA: antimycin A; df: degrees of freedom.

### Immune landscape of healthy human kidney

Despite the relative paucity of immune cells in healthy human kidney, we examined kidney-resident immune cells to delineate their steady-state phenotypes and functions. Sub-clustering of immune cells yielded 12 clusters (**Fig. 4a**). T cells (*CD3E^+^),* Natural Killer (NK) cells (*NKG7^+^CD3E^-^*), and a small B cell population (*CD79A*^+^) mainly expressing the immunoglobulin chain *IGHM* were identified (**Fig. 4b, Supplementary Fig. 7a**). Plasma cells (*CD38*^+^*XBP1*^+^) were scarce in healthy kidney tissue (**Supplementary Fig. 7b**). Myeloid clusters (*CD68*^+^) (**Fig. 4b)** displayed enrichment of phagocyte-related pathways including “receptor-mediated endocytosis”, “regulation of TLR signaling”, and “antigen processing and presentation via MHC class II” (**Supplementary Fig. 7c**).

**Figure 4.**
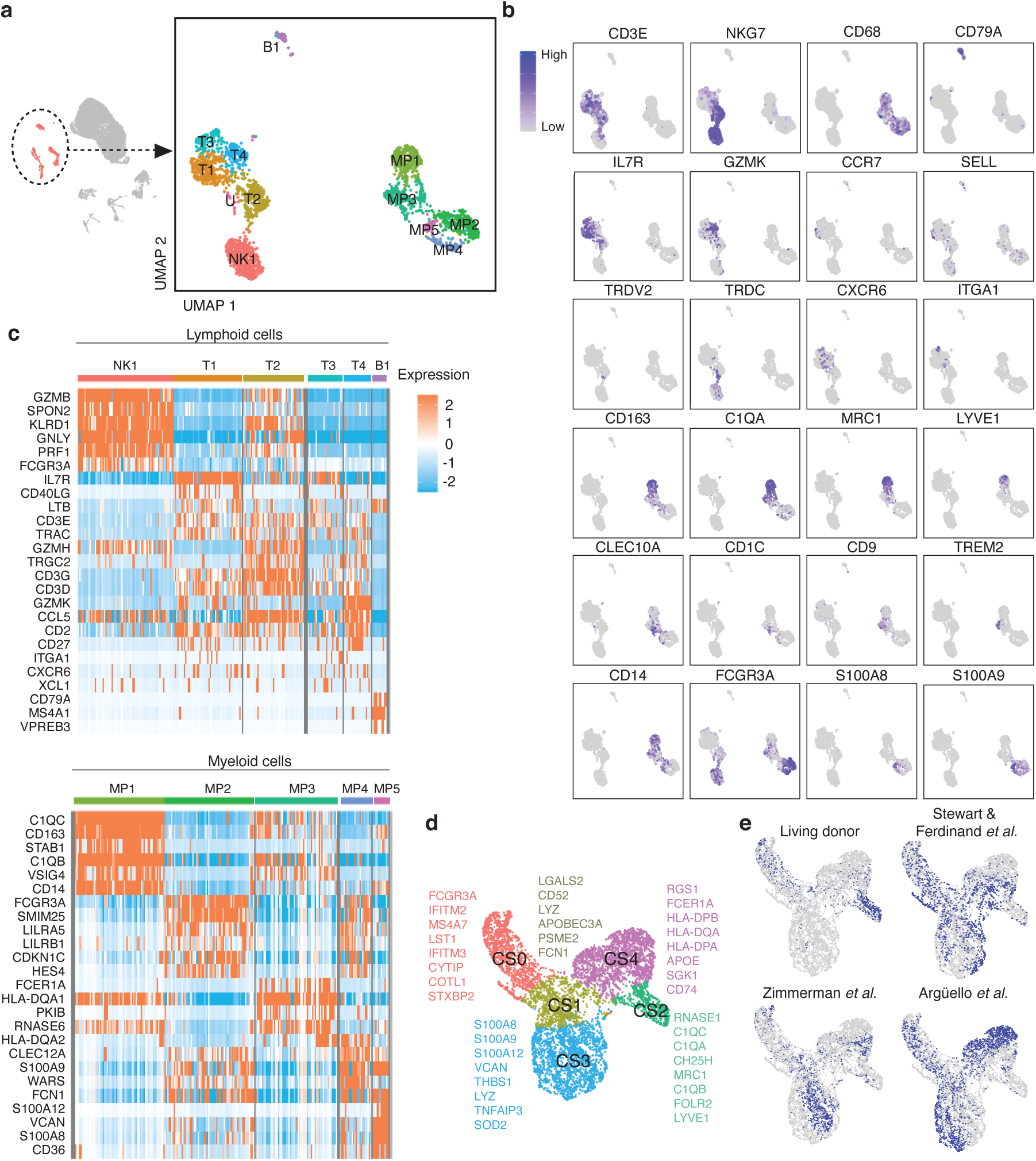
Identification and annotation of kidney immune cells. (**a**) Compartment-specific analysis of 2491 immune cells comprising 12 clusters and (**b**) cell type markers used for cluster annotations (**c**) Heatmap of cell-type defining and highly expressed genes by each cluster separated by lymphoid and myeloid lineage. (**d**) UMAP plot showing the living donor myeloid cell data clustered together with the same three published datasets to define five cell states across datasets and their respective cluster markers. (**e**) UMAP plots highlighting the distribution of dataset membership across the cell states.

T cell cluster T1 expressed CD4^+^ T helper (Th) cell genes (*IL7R*^+^*CD40LG*^+^*LTB* ^+^) and enrichment of “T-helper cell differentiation” and “Interleukin-7-mediated signaling” pathways (**Fig. 4c, Supplementary Fig. 7c**). T1 also included *CCR7^+^ SELL^+^* cells, suggesting central memory T cell identity (**Supplementary Fig. 8**)^36^. T2 demonstrates expression of a cytotoxic program (*GZMA, GZMB, GZMH, GNLY, PRF1*) alongside NK receptor genes (*KLRD1*, *KLRG1*), consistent with effector memory T cell or NKT cell identity (**Fig. 4c**), T2 also contained some gamma-delta (*γδ*) T cells, marked by co-expression of TCR chain components *TRDV2* and *TRDC* (**Fig. 4b**). T3 had sparse expression of resident memory T cell (Trm) markers (*CXCR6*, *ITGA1*), while T4 was marked by high *GZMK* expression, a marker of circulating age-associated memory T cells (**Fig. 4b, c**)^37^. FOXP3^+^CD4^+^ regulatory T cells were notably absent from scRNAseq and flow cytometry analyses (**Supplementary Fig. 9a**), while being observed in kidney pathologies^38, 39^, indicating they are likely recruited during inflammation. NK cell cluster NK1 displayed a cytotoxic gene program and broad *FCGR3A*(CD16) expression. Flow cytometry confirmed ∼95% of renal NK cells are CD56^dim^CD16^+^(**Supplementary Fig. 10a**). Low abundance of ILC2s, ILC3s and CD56^bright^ NK cells was suggested by a predictive classifier and confirmed by flow cytometry (**Supplementary Fig. 10b, c**).

As we noted differences in our lymphocytes signatures to those reported using other tissue sources, we directly compared lymphocytes in living donor kidney with tumor-unaffected renal tissue. We confirmed the presence of many similar immune populations across tissue sources, yet also observed differences in abundance and transcriptional signatures. When T cell and NK cell clusters were compared between these different tissue sources, alterations in checkpoint molecule expression (i.e *TIGIT*, *CTLA4*, *PDCD1*) were noted, with some of these differences also being observed at the protein level. We also observed high donor heterogeneity in immune infiltration and generally a greater proportion of immune cells in nephrectomy specimens, supporting the immune niche can be altered from healthy kidney.

Mononuclear phagocytes (MP) acquire tissue-adapted phenotypes and functions^40^. Definitively attributing macrophage or DC identity to myeloid populations based on gene expression alone is particularly challenging within the kidney due to a lack of consensus on lineage defining markers^41^ and here they are annotated more generally as five MP populations. Cluster MP1 highly expressed complement components (*C1QA*, *C1QB*, *C1QC*) and markers of alternative macrophage activation or anti-inflammatory function (*CD163*, *LYVE1*, *STAB1*, *MRC1*, *VSIG4*, *FOLR2*) (**Fig. 4b-c, Supplementary Fig. 12**). Efferocytosis receptor *MERTK* expression supports homeostasis or repair functions (**Supplementary Fig. 12**). MP3 contained cells expressing cDC2 markers (*CLEC10A*, *CD1C*), alongside a subgroup of cells co-expressing lipid-associated genes (*CD9*, *TREM2*, *APOE, APOC1*) (**Fig. 4b**). Similar populations have been identified as kidney-resident macrophages and are expanded in fibrotic tissues^42^. MP2 and MP4 (*FCGR3A*^+^*SIGLEC10*^+^*FCN1*^+^) resemble CD16^+^ non-classical monocytes (**Fig. 4b-c, Supplementary Fig. 12**). MP4 had elevated expression of *IL1B*, MHC Class-II genes, and *CX3CR1* while MP2 had higher expression of *CXCR4* and *FPR1* (**Supplementary Fig. 7d**). MP5 expressed markers of classical CD14^+^ monocytes (*S100A8*, *S100A9*, *CD14*, *VCAN*), yet was predominantly from one individual with elevated hemoglobin transcripts, indicative of increased circulating cells in this particular sample (**Fig. 4b, c, Supplementary Fig. 7e)**. Flow cytometry confirmed greater abundance of CD16^+^ cells in kidney relative to blood, as well as low proportions of CD14^+^CD16^-^ MPs resembling MP5 and the presence of MRC1^+^HLA-DR^+^ MPs in kidney that align with MP1 (**Supplementary Fig. 10e, f**).

### Identification of a distinct resident macrophage population in healthy kidney

Due to unique aspects of our study, including short ischemic times to which resident macrophages are especially sensitive^43, 44^, and use of flushed living donor-derived kidney tissue, we examined shared and unique MP populations in healthy kidney compared to those reported previously in kidney tissue from other sources. *CD68*^+^ cells from three prior studies^3, 45, 46^ were classified to match cluster identities of our study. MPs from these studies most resembled MP5 (classical CD14^+^ monocyte-like), the lowest abundance MP cluster in living donor samples (**Supplementary Fig. 13a**). MP3 (DC-like and lipid-associated MPs) as well as MP2 and MP4 (CD16^+^ non-classical monocyte-like) were shared across datasets. Strikingly, few cells from these studies corresponded to MP1 (resident macrophages) – the largest MP population in living donor kidney. Next, *CD68*^+^ cells from these prior studies^3, 45, 46^ and our study were merged, identifying five myeloid cell states (CS) across all studies (**Fig. 3d**). Based on transcriptomic profiles, CS2 and CS4 include resident macrophages and antigen-presenting cells, CS0 is consistent with non-classical CD16^+^ monocytes, CS3 represents classical CD14^+^ monocytes and CS1 may represent a transition state, supported by trajectory analysis (**Supplementary Fig. 13b, c**). CS2, which was almost entirely comprised of living donor kidney cells (**Supplementary Fig. 13b**), is defined by expression of genes associated with alternatively activated macrophages (*C1QA/B/C*^+^*RNASE1*^+^*CD163*^+^*LYVE1*^+^*FOLR2*^+^), in contrast to all other CS which expressed markers associated with monocytes and classically activated macrophages (S100 family members, *FCN1*, *LYZ*, and pro-inflammatory *SOD2*) (**Fig. 4d, Supplementary Fig. 13b**). CS2 constitutes the predominant MP population in healthy kidney (MP1), while CS3 and CS4 abundance is limited (**Fig. 4e)**

### Kidney-resident lymphocytes are antigen-experienced with distinct gene expression

Due to unexpected heterogeneity and novel transcriptional profiles in kidney lymphocyte populations (**Fig. 4a-c**), we directly compared lymphocyte proportions, signatures, and phenotypes to those in healthy donor blood. Increased proportions of NK (CD3^-^CD56^+^) and NKT cells (CD3^+^CD56^+^) were noted in kidney, while T cell (CD3^+^CD56^-^) abundance was unchanged (**Fig. 5a**). CD8^+^ T cells were present in higher proportions than CD4^+^ T cells in kidney and the presence of *γδ*T cells was validated by flow cytometry (**Fig. 5b, Supplementary Fig. 9a**).

**Figure 5:**
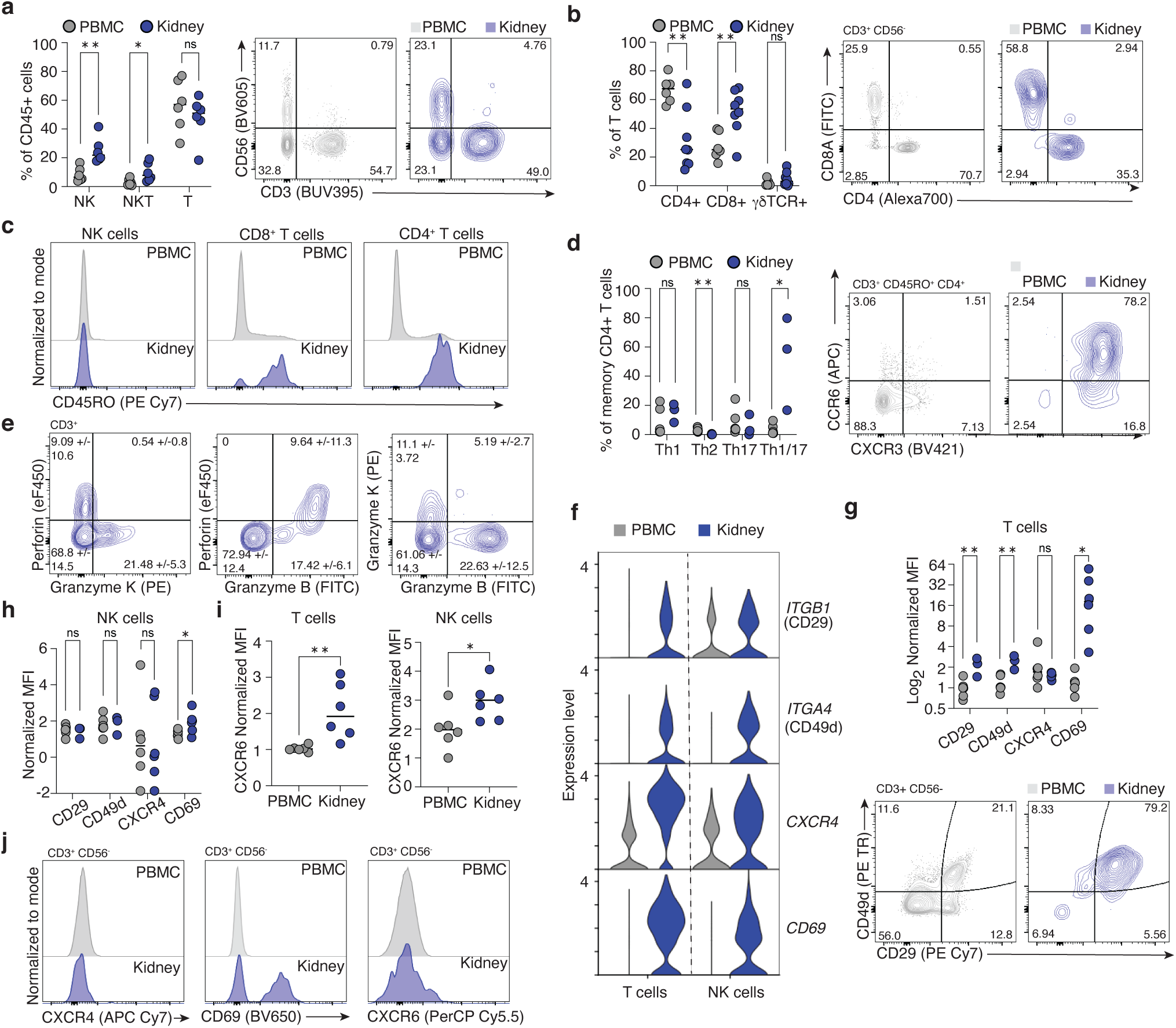
Characterization of kidney-resident T and NK cells. (**a**) NK cells (p=0.0025, t=3.998, df=10) and NKT cells (p=0.0327, t=2.476, df=10) are proportionally enriched in kidney relative to blood, while T cell (p=0.379, t=0.918, df=10) abundance is unchanged. (n= 6) (**b**) Within the kidney T cell population, there is an enrichment of CD8^+^ T cells (p=0.0060, t=3.327, df=12) and a reduction in CD4+ T cell (p=0.0025, t=3.815, df=12) abundance with no change in TCRgd^+^ T cells (p=0.2158, u=14) relative to blood. (n=6) (**c**) Kidney T cells are predominantly antigen-experienced, marked by expression of CD45RO, while NK cells express minimal CD45RO. (**d**) Within kidney memory CD4^+^ T cells, there is an enrichment in the Th1/17 subpopulation (CXCR3^+^CCR6^+^) (p=0.0238, u=0) and a reduction in Th2 subpopulation (CRTh2^+^) (p=0.0098, t=3.513, df=7) abundance relative to blood while Th1 (CXCR3^+^) (p=0.3810, u=5) and Th17 (CCR6^+^) (p=0.5476, u=6) proportions were unchanged. (n=3) (**e**) T cells expressing Granzyme K do not co-express perforin, indicating that they are a distinct T cell subset from Granzyme B^+^Perforin^+^ cytotoxic T cells. (**f**) Violin plots showing differential gene expression of select markers in kidney T cells and NK cells relative to blood. (**g**) Surface levels of CD29 (p=0.0061, t=3.869, df=7), CD49d (p=0.0027, t=4.519, df=7) and CD69 (p=0.0203, t=2.756, df=10) were higher on kidney T cells relative to blood as measured by flow cytometry, while CXCR4 (p=0.5887, u=14) was not (n=6). (**h**) Surface levels of CD69 (p=0.0427, t=2.321, df=10) was higher on kidney NK cells relative to blood while CD29 (p=0.6899, t=0.4159, df=7), CD49d (p=0.9040, t=0.1250, df=7), and CXCR4 (p=0.9326, t=0.0868, df=10) were not. (n=6) (**i**) CXCR6 abundance was higher at the protein level on both T cells (p=0.0086, t=3.258, df=10) and NK cells (p=p=0.0364, t=2.414, df=10) relative to blood. (**j**) Histograms showing no difference in CXCR4, increased CD69 and increased CXCR6 protein abundance in kidney T cells relative to blood. Group-to-group differences were assessed using two-tailed unpaired T tests for variables following a normal distribution, and Mann-Whitney tests for variables with a non-parametric distribution., *p<0.05;**p<0.01;***p<0.001;****p<0.0001.

To identify specific markers and transcriptional profiles of kidney-resident lymphocytes, we integrated our dataset with public PBMC scRNA-seq datasets^47, 48^(**Supplementary Fig. 14a-j, Supplementary Table 4**) and validated differences by flow cytometry. Unsurprisingly, blood lymphocytes exhibited higher expression of naïve T cell genes (*CCR7*, *SELL*, *LEF1*, *TCF7*). In contrast, the tissue residency-associated transcription factor *PRDM1* (BLIMP-1)^49^ was upregulated in kidney lymphocytes, as was *CD69,* which marks Trms in several organs and prevents tissue egress via S1PR1 antagonism^50^ (**Supplementary Fig. 14j**). Antigen-experienced T cells upregulate CD45RO and can become Trm^51^. 60-98% of kidney CD4^+^ and CD8^+^ T cells were CD45RO^+^ in contrast to low proportions of memory T cells in blood (**Fig. 5c**). NK cells with memory functions may also express CD45RO^52^; however, this was not observed in renal NK cells (**Fig. 5c**). Flow cytometry confirmed elevated CD69 on T cells and NK cells, with CD69-CD103 co-expression by CD8^+^ T cells, consistent with a Trm phenotype (**Supplementary Fig. 9c**). Further characterization of memory CD4^+^ T helper (Th) cell subsets revealed enrichment of Th1/17 cells with reduced Th2 marker expression (**Fig. 5d, Supplementary Fig. 9b**).

We also sought to validate Granzyme K production in kidney lymphocytes, as T4 cluster was marked by high *GZMK* expression. In agreement with scRNAseq findings, Granzyme K was detected in 21% of kidney T cells (**Fig. 5e**), with minimal co-expression with Granzyme B, indicating that Granzyme K^+^ T cells form a distinct subset of renal T cells (**Fig. 5e**). Most Granzyme K^+^ T cells also did not have detectable perforin expression (**Fig. 5e**), in line with Granzyme K produced by these T cells having extracellular functions rather than the canonical cytolytic function of granzymes dependent on intracellular delivery via perforin.

Kidney lymphocytes were distinguished from circulating lymphocytes by elevated expression of chemokine receptors (*CXCR4*, *CXCR6*), integrin components (*ITGB1*, *ITGA4*), and inhibitory NK receptors (*KLRD1*, *KLRC1*) (**Fig. 5f, Supplementary Fig. 14j**). Flow cytometry confirmed VLA-4 integrin components *α*4 (CD49d) and *β*1 (CD29) were highly expressed in renal T cells suggesting VLA-4 contributes to their residency or function (**Fig. 5g**). This is consistent with expression of VLA-4 ligands fibronectin and VCAM-1 in kidney^53^. Kidney NK cells have higher levels of CD69 compared to circulating NK cells, while no difference in CD29 or CD49d was detected (**Fig. 5h**). Finally, CXCR6 protein expression was elevated on kidney T and NK cells, while CXCR4 was not, despite high gene expression (**Fig. 5g, h, i**). Notably, renal myeloid cells expressed *CXCL16*, the chemokine ligand for CXCR6, indicating participation in lymphocyte recruitment, supported by significant aggregate rank scores using cell-cell communication inference (**Supplementary Fig. 9d**, **Supplementary Tables 5, 6**).

Other differentially expressed genes suggest tissue-adapted function of kidney lymphocytes. *AREG,* encoding the growth factor amphiregulin, was highly expressed by NK1 and validated by flow cytometry (**Supplementary Fig. 10d, Supplementary Fig, 11f, g**), suggesting tissue-reparative functions. The prostaglandin E2 (PGE2) receptor *PTGER4* and prostaglandin D synthase *PTGDS* were upregulated (**Supplementary Fig. 14j**), indicating kidney lymphocytes synthesize and recognize prostaglandins, known mediators of kidney function^54^. PGE2 promotes Th17 and Th1/17 cell development and function, perhaps explaining Th1/17 cell enrichment in kidney (**Fig. 4d**)^55^. Collectively these studies capture the heterogeneity of myeloid and lymphocyte populations within healthy human kidney and provide an important reference of immune cell phenotypes and functions at steady state.

## Discussion

We present a scRNAseq atlas of healthy human kidney using biopsies from living donors. Our resolution of healthy kidney PT, endothelial, epithelial, and immune subpopulations will inform future studies addressing underling mechanisms of kidney pathologies, including chronic kidney disease, fibrosis, IRI, renal cancer and allograft rejection.

The sex-balanced design in the present study enabled novel examination of sex-based dichotomy in gene expression among human kidney cell populations. Prior studies were constrained by small sample size and use of animal models, or instead used bulk transcriptional analysis where sex-specific signatures of individual kidney cell populations cannot be resolved^19, 23, 57^. Our study is aligned with the conclusion of scRNAseq studies in mouse by Ransick *et al.* ^23^ that PT cells are sexually dimorphic. However, the overlap in sexually dimorphic PT genes between human and mouse is small, perhaps due to distinct orthologues in mouse, small number of samples sequenced, or true biological differences between human and mouse.

We report striking sex-based transcriptional differences in PT cells, suggesting higher baseline metabolic activity in males, and enhanced expression of antioxidant genes in females. We validated these sex-based observations at the level of gene expression, metabolite generation, and metabolic function *in vitro*. Increased oxidative stress is reported in males^58^, while female sex hormones augment antioxidant gene transcription^59^. Metallothionein genes (*MT1F, MT1G, MT1H*), which are potent endogenous antioxidants^60^, were increased in female PT cells. Metallothionein depletion exacerbates diabetic and hypoxia-induced kidney injury^61, 62^, whereas augmented expression is protective^63^. Several sex-altered genes further relate to cysteine-glutathione metabolism. Glutathione is critical to cellular antioxidant defences^64^ and glutathione metabolism exhibits sexual dimorphism^22, 65^. These sex-based differences in PT gene expression discovered by use of scRNAseq which can capture transcripts localized to the mitochondria and cytosol, may provide insights into the well-recognized, but previously unexplained sexual dimorphism observed in most kidney diseases. In particular, why females may be less susceptible to metabolism-related kidney injury ^6–8, 66, 67^.

Our study provides a steady-state map of the kidney immune niche. Kidney T cells are predominantly Trms and exhibit unique phenotypes previously unreported in kidney, including Granzyme K^+^ T cells. The function of Granzyme K^+^ T cells in humans is poorly characterized, and here we show that Granzyme K^+^ T cells are a distinct subset separate from Granzyme B^+^Perforin^+^ T cells in the kidney. The lack of perforin co-expression suggests that Granzyme K produced by renal T cells may have extracellular targets, such as inducing endothelial cell activation^68^, promoting sensitivity to LPS-induced inflammation^69^, and regulating angiogenesis^70^.

Renal CD4^+^ memory Th cells are skewed towards a Th1/17 phenotype, which may be relevant to Th17-related kidney diseases including glomerulonephritis, lupus nephritis, and transplant rejection^71, 72^. Renal abundance of CD56^+^CD16^+^ NK cells with high expression of amphiregulin compared to circulating NK cells suggests non-canonical tissue-adapted functions. We demonstrate an enrichment of a resident macrophage population with little-to-no presence in prior datasets from discarded deceased donor or tumor nephrectomy specimens, suggesting altered kidney environments impact this myeloid population. Indeed, sensitivity of self-renewing resident macrophage populations to extended ischemic injury and inflammation is reported^44^. Additional comparison of lymphocyte populations in tumor-unaffected versus living donor renal tissue revealed alterations in tumor-unaffected tissue relative to the steady-state immune niche in healthy living donor kidney. Increased B and T cell proportions, increased expression of activation and exhaustion-associated molecules by lymphocytes, in addition to a trend for increased immune infiltration in nephrectomy specimens was observed (**Supplementary Fig. 11**), in agreement with prior reports that tumour-affected kidneys can have altered immune infiltrates^5, 56^. Future studies exploring alterations in immune cells in tumor-unaffected kidney tissue of renal cancer patients may have implications for development of immunotherapies.

Collectively, our description of healthy human kidney provides a reference point for understanding the cellular basis of kidney disease development, represents a ‘normal’ target for stem cell-derived kidney organoids, and expands our understanding of the complexity of sex-based gene expression and kidney-resident immune populations.

## Abbreviations

AUC: Area under the curve
CCD: Cortical collecting duct
CNT: Connecting tubule
CTAL: Cortical thick ascending limb of the loop of Henle
DC: Dendritic cell
DCT: Distal convoluted tubule
Endo: Endothelial
IC-A: Intercalated cells type A
IC-B: Intercalated cells type B
IRI: Ischemia-reperfusion injury
LogFC: Log Fold Change
MP: Mononuclear phagocyte
MHC: Major histocompatibility complex
NK cell: Natural killer cell
Non-PT: Non proximal tubular parenchymal cell
PBMCs: Peripheral blood mononuclear cells
PGE2: Prostaglandin E2
PT: Proximal tubule
RBC: Red blood cell
scRNAseq: Single cell RNA sequencing
STC: Scattered tubular cell
TCA: Tricarboxylic Acid
TCR: T cell receptor

## Acknowledgments

First and foremost, we would like to thank the kidney transplant patients who made this work possible. We would also like to thank the nurses, physicians and surgeons at Toronto General Hospital and the Ajmera Transplant Centre Biobank for efforts to obtain tissue samples, and acknowledge technical support provided by the Princess Margaret Genomics Centre, particularly Troy Ketela and Julissa Tsao, and the Princess Margaret Cancer Centre flow cytometry core. CMM was supported by the Menkes Family Fellowship and an Ajmera Transplant Centre fellowship. JMM was supported a QE II/Aventis Pasteur Graduate Scholarship and a Peterborough K.M. Hunter Foundation scholarship. This work supported by funding from the Canadian Institutes for Health Research (CIHR) grant 168960 to SQC and AK and the Ajmera Transplant Centre/Toronto General and Western Hospital Foundation (TGWHF). SQC was also supported by the Medicine by Design program (Canada First Research Excellence Fund) and Canada Foundation for Innovation (CFI) grant 38308. AK was supported by Kidney Foundation of Canada (KFOC) Predictive Biomarker grant KFOC160010, CIHR 347479, CFI grant 37205, KFOC Biomedical Research grant KFOC160010, and Kidney Research Scientist Core Education and National Training (KRESCENT) program grants CIHR148204, KRES160004, and KRES160005, as well as funding from TGWHF (TGTWF 1617-464; TGTWF MKFTR 1718-1268).

## Author contributions

CMM, JMM, AK, and SQC designed and implemented the study. AK and SS established the ATC biobank and the infrastructure required for sample retrieval. SS, CMM, JMM, SZ, KAL, AF and AK liaised with patient care teams to obtain tissues for study. CMM, JMM, JAM, SCF, JA, AK and SQC developed tissue dissociation protocols with input from LL and SAM, and performed experiments. SCF, OZ, HR, RA, contributed to PT validation experiments. CMM, JMM and LZ designed and implemented the bioinformatic pipeline with advice and assistance from MK, DP, BW, GDB, AK and SQC. CMM, JMM, SAM, AK and SQC provided tissue and immune compartment expertise and annotated cell types. BW, SAM, GDB, AK, and SQC supervised the work. CMM, JMM, AK, and SQC wrote the manuscript, which all authors reviewed and edited.

## Competing interests

Authors declare that they have no competing interests

## Materials and correspondence

Correspondence and requests for materials should be addressed to Dr. Sarah Crome and Dr. Ana Konvalinka

## Materials and Methods

### EXPERIMENTAL MODEL AND SUBJECT DETAILS

#### Human Specimens

Kidney tissue from tumour-unaffected nephrectomy specimens was used for initial method optimization. Pre-implantation core biopsies were obtained from living donor kidneys after organ retrieval and flushing. 20 live donor kidney samples (10 male donors and 10 female donors) were processed for sequencing. Additional living donor kidney samples were used in flow cytometry experiments for method optimization and immunophenotyping. All experiments were conducted with institutional ethics approval from University Health Network (CAPCR: 18-5914.0, Living donor; CAPCR: 18-5489.0, Tumour nephrectomy). Patient demographic information for sequenced samples is summarized in **Supplementary Table 7**. All patients provided informed written consent for inclusion in this study.

#### Murine Specimens

Murine kidneys from C57BL/6 mice (AUP: 6156) were used for digestion optimization experiments.

### EXPERIMENTAL METHOD DETAILS

#### Tissue digestion and CD45-enrichment

All living donor samples used for sequencing were processed within one hour of organ retrieval. Briefly, biopsies were collected in RPMI 1640 (Gibco, cat # 11875119) on ice, and mechanically dissociated with a blade before enzymatic digestion at 37°C with 0.1 mg/ml DNase I (STEMCELL, cat # 07470), 3300 CDA units/ml Collagenase MA (VitaCyte, cat # 001-2030) and 1430 NP units/ml BP neutral protease (VitaCyte, cat # 003-1000) for 20 minutes at 37°C with intermittent agitation in an dissociation protocol optimized to maximize viability and to preserve representation of rare and fragile cell populations (**Supplementary Fig. 15**). Cell suspensions were filtered through 35µm cell strainer snap-cap FACS tubes (Falcon, cat# 352235) and a plunger from a 1ml syringe was used to gently mash remaining tissue in the strainer before rinsing strainer lid with 1:1 volume of FBS (HyClone, cat # SH3039603PM) on ice. A low frequency (<1%) of immune cells in the single cell suspension from a kidney biopsy core (**Fig. 1a**) necessitated immune enrichment in 10 samples (5 males, 5 females) using magnetic EasySep Human CD45 depletion kit II (STEMCELL, cat# 17898), as per the manufacturer’s modified instructions for positive selection of CD45-expressing cells.

#### Single-cell RNA sequencing

Samples were prepared according to 10X Genomics Single Cell 3’ v3 Reagent kit user guide^73^. The pilot sequencing sample from nephrectomy tissue was sequenced using 10X Genomics Single Cell 5’ v2 Reagents. Samples were washed twice in PBS (Life Technologies) plus 0.04% BSA, and viability was determined by a hemocytometer (Thermo Fisher) via Trypan Blue staining. Following counting, the appropriate volume for each sample was calculated for a target capture of 9,000 cells. For CD45-enriched samples, all cells were sequenced. Samples that were too low in cell concentration as defined by the user guide were washed, re-suspended in a reduced volume and counted again using a haemocytometer prior to loading onto the 10x single cell B chip. After droplet generation, samples were transferred onto a pre-chilled 96 well plate (Eppendorf), heat sealed and incubated overnight in a Veriti 96-well thermos cycler (Thermo Fisher). The next day, sample cDNA was recovered using Recovery Agent provided by 10x and subsequently cleaned up using a Silane DynaBead (Thermo Fisher) mix as outlined by the user guide. Purified cDNA was amplified for 11 cycles before being cleaned up using SPRIselect beads (Beckman). Samples were diluted 4:1 (elution buffer (Qiagen):cDNA) and run on a Bioanalyzer (Agilent Technologies) to determine cDNA concentration. cDNA libraries were prepared as outlined by the Single Cell 3’ Reagent Kits v3 user guide with modifications to the PCR cycles based on the calculated cDNA concentration.

The molarity of each library was calculated based on library size as measured bioanalyzer (Agilent Technologies) and qPCR amplification data (Roche). Samples were pooled and normalized to 1.5 nM. Library pool was denatured using 0.2N NaOH (Sigma) for 8 minutes at room temperature, neutralized with 400mM Tris-HCL (Sigma). Library pool at a final concentration of 300pM were loaded to sequence on Novaseq 6000 (Illumina). Samples were sequenced with the following run parameters: Read 1-28 cycles, Read 2-90, index 1-10 cycles, index 2-10 cycles. Across samples, cells were sequenced to a target depth of 40,000 reads per cell. Mapping and quantification were performed using the 10X Genomics CellRanger pipeline version 3.1.0. Cell metric summaries for each sample in **Supplementary Table 8**.

#### Single-nucleus RNA sequencing

A pilot single-nucleus RNA sequencing experiment was undertaken to compare single cell versus single nuclear results from a matched sample. The biopsy was collected fresh and divided into 8 segments, evenly distributed to be processed fresh for single cell RNA sequencing as above, and the remainder was flash frozen in liquid nitrogen. The sample was later retrieved from liquid nitrogen and processed on dry ice according to the protocol in^74^ with a lysis buffer containing: 0.32 mM sucrose (BioShop SUC507.1), 5 mM CaCl2 (VWR, 97062-820), 3 mM MgCl2 (Thermo Fisher AM9530G), 20 mM Tris-HCl pH 7.5 (Thermo Fisher, 15567027), 0.1% TritonX-100 (Sigma Aldrich T8787-50ML), 0.1 mM EDTA pH 8.0 (Thermo Fisher AM9260G), 40 U/ml Protector RNAse inhibitor (Sigma Aldrich 3335399001) in UltraPure DNAse/RNAse-free water (Thermo Fisher 10977015). The nuclei were captured and sequenced using 10X Genomics Single Cell 3’ v3 Reagents as above.

#### Data quality control, clustering, differential expression, pathway analysis and cell-cell interaction inference

Original study recruitment included samples from 20 donors, however, data from one male donor was poor quality and was excluded from downstream analysis. Thus, our final dataset consisted of 19 donors (10 female, 9 male), with 10 CD45-enriched samples (5 female, 5 male) and 9 samples not enriched for CD45^+^ cells referred to as “total kidney” (5 female, 4 males). To preserve representation of rare cell types with uniquely expressed genes, we retained genes expressed in a minimum of 1 cell in the individual datasets.

Ambient RNA contamination was corrected using the AutoEst function in SoupX^75^ (**Supplementary Fig. 16**). DoubletFinder^76^ was used to identify and remove cells most likely to be doublets, rather than implementation of a maximum gene or feature threshold. For total samples, a high doublet rate threshold of 7.5% was applied (as utilized in comparable studies^77^), while for CD45-enriched samples, the doublet rate was calculated as 0.8% per 1000 cells captured, as per 10X Genomics estimated doublet rates^73^. The individual datasets were then merged. Upon merging all of the individual datasets, the cells clustered according to cell type rather than donor/batch, and importantly, no batch correction of the data was required.

Cell type-specific thresholds were set to remove low quality cells. For immune cells (clusters expressing *PTPRC*), all cells with >10% of UMIs mapped to mitochondrial genes were removed, along with cells that had low transcript abundance (<1000) or gene diversity (<200 unique genes). Separately, prior to removing cells with low transcripts/features, data was mined for the presence of granulocyte lineage cells such as neutrophils which are often removed by typical QC thresholds due to high RNAse activity and low gene content, however very few neutrophils (>20) were identified by marker expression in the raw data across all samples. For parenchymal cells, all cells with >40% of mitochondrial-mapped UMIs were removed; this high threshold was imposed due to known high mitochondrial content of proximal tubular cells^78^. Additionally, cells with low transcript abundance <1000) and low gene diversity (<750 unique genes) were removed. Cells expressing hemoglobin genes (*HBB*, *HBA1/2*) (n=160) were removed. Following normalization (SCTransform^79^) and feature selection (M3Drop/DANB^80^), principal component analysis was used for dimensionality reduction (RunPCA) and cells were clustered using the Louvain algorithm with 30 principal components (FindNeighbors and FindClusters) (Seurat^81^). Clusters were visualized using UMAP algorithm^82^.

The dataset was divided into 3 broad subgroups identified as being Immune (*PTPRC*^+^) or Parenchymal (Proximal Tubular (expressing *CUBN*, *HNF4A*, *SLC34A1*, *LRP2*, *SLC17A1*) or non-Proximal Tubular) in origin. These subgroups were re-clustered and further annotated using a curated marker list (**Supplementary Table 6**). Cluster defining genes were identified by Seurat’s FindMarkers^81^.

Ranked gene lists were generated using Wilcoxon rank sum testing from the presto package (wilcoxauc function)^83^ were used as input for pathway analysis using GSEA^84^. Reference gene sets were acquired from the Bader lab repository (http://download.baderlab.org/EM_Genesets/) – Geneset used: (Human_GOBP_AllPathways_no_GO_iea_January_13_2021_symbol.gmt.txt). To identify pathways enriched in immune cell clusters, the ranked gene lists were generated for each cluster comparing that cluster versus all other clusters.

Cell-cell communication was inferred from the sequencing data using LIANA which generates a consensus ranking across several methods^85^. The OmniPath interaction database was used^86^ with the following methods for inferring interactions implemented through the package: SingleCellSignalR^87^, iTalk^88^, NATMI^89^, Connectome^90^, CellChat^91^ and CellPhoneDB^92^. Results are summarized in **Supplementary Table 5**. Separately, SingleCellSignalR, NATMI, iTALK and Connectome methods were used to generate a consensus score using the CellPhoneDB database to infer interactions inclusive of multimeric complexes as accounted for in the CellPhoneDB interaction database, summarized in **Supplementary Table 6**.

#### Identification of innate lymphoid cells

A predictive tool for cell type classification (scPred^93^) was trained on single-cell data generated from flow cytometry-sorted ILCs^94^ and T cells^95^. Using this classifier, some cells present within our dataset were putatively identified as ILCs.

#### Transcription factor analysis

Top cluster defining genes for PT5 and PT3, respectively were uploaded to CHEA3^96^ (https://maayanlab.cloud/chea3/), and the top 10 predicted upstream regulators were identified.

#### Comparison of kidney immune cells to PBMCs

To identify differences in gene expression between T cells and NK cells from peripheral blood versus kidney, PBMC data (GSE148665)^47^ was integrated with the immune only kidney data using Harmony^97^. A second independent PBMC dataset^48^, was separately integrated with the kidney data for dataset-independent validation. NK cells and T cells (clusters expressing *NKG7* and/or *CD3E*) were compared using Seurat’s FindAllMarkers function. Violin plots and volcano plots were created using Seurat and EnhancedVolcano^98^.

#### Comparison of Myeloid cells

To identify differences in myeloid cell populations in living kidney donors compared to publicly available human kidney single-cell RNA sequencing datasets from tumour nephrectomy or deceased donor tissue sources, CD68-expressing clusters from Stewart & Ferdinand *et al*.,^3^ Zimmerman *et al*.,^45^ and Argüello *et al*.^46^ were scored using a random forest classifier (SingleCellNet^99^) to identify cells from the published datasets corresponding to the five myeloid clusters in the living donor data. Separately, all myeloid cells from this data and the three published studies were integrated and clustered to identify cell states using OCAT^100^. The datasets were also integrated and batch corrected using Seurat v3 integration (FindIntegrationAnchors and IntegrateData functions). The cell state identities from OCAT were mapped onto the integrated object and marker genes of cell states were identified using Seurat’s FindAllMarkers function. Lineage analysis by pseudotime inference was applied to the OCAT-identified clustering of the combined myeloid populations using slingshot^101^, without indicating any clusters as either start or end points.

#### Sex differences analysis

Principal component analysis (PCA) followed by Varimax rotation was performed on all major parenchymal and immune populations. Varimax-rotated principal components 2:25 were serially plotted against component 1, to identify whether a separation on the basis of sex was evident. If seen, the top 100 genes (50 from each end of the gene loading list) associated with the Varimax-rotated principal component were retained for further analysis.

Sex differences in proximal tubular cells were identified using sparse partial least squares discriminant analysis (sPLS-DA) in mixOmics^102^. Using the tuning function (tune.splsda), the optimal values for sparsity parameters were determined to be 1 component with 80 variables (genes). To test the classifier, the data were separated into a training dataset (⅔ of cells sampled) and a query dataset (remaining ⅓). Next, our 80-gene signature was applied to an external dataset (Liao *et al*.^20^) for validation. Here, the entire living donor dataset was used as the training dataset and the external dataset was used as the query dataset. To determine the contribution of sex chromosome encoded genes to the model, all X- and Y-chromosome encoded genes were removed from the datasets prior to analysis, where the tuned parameters identified the optimal model to include 1 component with 15 variables. This 15-gene signature was also validated in the Liao *et al*. dataset. Hierarchical structure, zero inflation, and pseudoreplication bias in single-cell data pose specific challenges for differential expression analyses^103–105^. To circumvent these limitations, we implemented a mixed effects model using MAST^105, 106^. For differential expression testing between male and female proximal tubule cells, the dataset was filtered to include only genes which were expressed in each sample (9792 genes). Differential expression testing was conducted using MAST with a random effect for sample (zlm∼ cellular detection rate + donor sex + (1| sampleID)). As this approach excluded genes expressed exclusively by one sex (e.g. Y chromosome encoded genes, and XIST), such genes were added to MAST differentially expressed genes (MAST+) for comparison with the results of the other methods (Varimax, sPLS-DA).

All significant genes returned using MAST analysis were subjected to enrichment analysis (GSEA^84, 107^) using reference gene sets acquired from the Bader lab repository: (http://download.baderlab.org/EM_Genesets/); Geneset used: (Human_GOBP_AllPathways_no_GO_iea_January_13_2021_symbol.gmt.txt).

#### Cryopreservation

Cells from additional (non-sequenced) fresh living donor biopsies or cells remaining following 10X cell capture for sequencing were resuspended in 90% human serum (Sigma, cat# H4522) and 10% DMSO for cryopreservation and cooled to -80°C in a Mr.Frosty (Sigma, cat #C1562), then transferred to liquid nitrogen for long term storage.

#### Flow Cytometry

After fresh tissue digestion, cells were washed in PBS + 2% FCS before staining. Cryopreserved cells were thawed and washed twice in PBS + 2% FCS. Cells were incubated at 4°C for 15 minutes with an Fc receptor blocker (BioLegend TruStain FcX, cat # 422302) according to manufacturer instructions before cocktails of surface antibodies were added for 30 minutes at 4°C. If intracellular targets/transcription factors were included in the panel, cells were resuspended in FOXP3 transcription factor fix perm buffer (eBio, cat # 00-5523-00) and stained with intracellular antibodies in 1X permeabilization buffer (eBio, cat # 00-8333-56). If no intracellular targets were included in the staining panel, cells were fixed in 2% PFA (Thermo Scientific, cat # J19443) after surface staining.

Cells were stained with the following surface antibodies: Anti-human CD8a FITC (1:100, clone RPA-T8, BioLegend, cat # 301050), Anti-human TCRgd FITC (1:100, clone B1, BioLegend, cat # 331208), Anti-human CD3 FITC (1:100, clone UCHT1, BioLegend, cat # 300440), Anti-human CD8a PerCP (1:50, clone RPA-T8, BioLegend, cat # 301030), Anti-human CXCR6 PerCP Cy5.5 (1:50, clone K041E5, BioLegend, cat # 356010), Anti-human CCR8 PE (1:100, clone L263G8, BioLegend, cat # 360604), Anti-human CD127 PE (1:50, clone hIL-7R-M21, BD Biosciences, cat # 557938), Anti-human CD15 PE (1:100, clone W6D3, BD Biosciences, cat # 562371), Anti-human CD163 PE (1:50, clone GHI/61, BioLegend, cat # 333606), Anti-human CD49d PE Dazzle 594 (1:100, clone 9F10, BioLegend, cat # 304325), Anti-human CRTh2 PE Dazzle 594 (1:50, clone BM16, BioLegend, cat # 350126), Anti-human CD31 PE Dazzle 594 (1:100, clone WM59, BioLegend, cat # 303130), Anti-human CD16 PE Dazzle 594 (1:100, clone 3G8, BioLegend, cat # 302054), Anti-human CD45 PE-CF594 (1:100, clone HI30, BD Biosciences, cat # 562279), Anti-human CD29 PE Cy7 (1:100, clone TS2/16, BioLegend, cat # 303025), Anti-human CD45RO PE Cy7 (1:50, clone UCHL1, BD Biosciences, cat # 560608), Anti-human MerTK PE Cy7 (1:50, clone 590H11G1E3, BioLegend, cat # 367610), Anti-human TIGIT PE Cy 7 (1:50, clone MBSA43, Invitrogen, cat # 25-9500-42), Anti-human CD94 APC (1:100, clone HP-3D9, eBioscience, cat # 17-5094-42), Anti-human CCR6 APC (1:25, clone G034E3, BioLegend, cat # 353416), Anti-human CD206 APC (1:50, clone 15-2, BioLegend, cat # 321110), Anti-human CD4 Alexa700 (1:50, clone RPA-T4, eBioscience, cat # 56-0049-42), Anti-human CD127 Alexa700 (1:50, clone eBioRDR5, eBioscience, cat # 56-1278-42), Anti-human CXCR4 APC Cy7 (1:50, clone 12G5, BioLegend, cat # 306528), Anti-human CTLA4 APC Cy7 (1:25, clone BNI3, BioLegend, cat # 369634), Anti-human CD56 APC Cy7 (1:50, clone HCD56, BioLegend, cat # 318332), Anti-human CD45 APC Cy7 (1:100, clone HI30, BioLegend, cat # 304014), Anti-human CD14 APC eF780 (1:100, clone 61D3, eBioscience, cat # 47-0149-42), Anti-human CXCR3 BV421 (1:50, clone G025H7, BioLegend, cat # 353716), Anti-human CD13 BV421 (1:50, clone WM15, BioLegend, cat # 301716), Anti-human TCRgd BV510 (1:100, clone B1, BioLegend, cat # 331220), Anti-human TCRab BV510 (1:100, clone IP26, BioLegend, cat # 306734), Anti-human CD5 BV510 (1:100, clone L17F12, BioLegend, cat # 364018), Anti-human FcER1 BV510 (1:100, clone AER-37, BioLegend, cat # 334626), Anti-human CD303 BV510 (1:100, clone 201A, BioLegend, cat # 354232), Anti-human CD123 BV510 (1:100, clone 6H6, BioLegend, cat # 306022), Anti-human CD34 BV510 (1:100, clone 581, BioLegend, cat #343528), Anti-human CD20 BV510 (1:100, clone 2H7, BioLegend, cat # 302340), Anti-human CD3 BV510 (1:100, clone OKT3, BioLegend, cat # 317332), Anti-human CD14 BV510 (1:100, clone M5E2, BioLegend, cat # 301842), Anti-human CD19 BV510 (1:100, clone HIB19, BioLegend, cat # 302242), Anti-human CD4 BV510 (1:100, clone RPA-T4, BioLegend, cat # 300546), Anti-human CD56 BV605 (1:50, clone HCD56, BioLegend, cat # 318334), Anti-human CD69 BV650 (1:100, clone FN50, BioLegend, cat # 310934), Anti-human CD8a BV650 (1:50, clone RPA-T8, BioLegend, cat # 301042), Anti-human CD326 BV650 (1:100, clone 9C4, BioLegend, cat # 324226), Anti-human CD107a BV750 (1:50, clone H4A3, BioLegend, cat # 328638), Anti-human CD103 BV711 (1:100, clone Ber-ACT8, BioLegend, cat # 350222), Anti-human CD10 BV711 (1:100, clone HI10a, BioLegend, cat # 312226), Anti-human CD45 BV711 (1:100, clone HI30, BioLegend, cat # 304050), Anti-human CD3 BV785 (1:100, clone OKT3, BioLegend, cat # 317330), Anti-human HLA-DR BV785 (1:50, clone L243, BioLegend, cat # 307642), Anti-human PD-1 BV785 (1:50, clone EH12.2H7, BioLegend, cat # 329930), Anti-human CD45 BUV395 (1:100, clone HI30, BD Biosciences, cat # 563792), Anti-human CD16 BUV395 (1:100, clone 3G8, BD Biosciences, cat # 563785), Anti-human CD3 BUV395 (1:100, clone UCHT1, BD Biosciences, cat # 563546), Anti-human CD69 BUV496 (1:50, clone FN50, BD Biosciences, cat # 750214), Anti-human CD16 BUV737 (1:100, clone 3G8, BD Biosciences, cat # 564434). The following antibodies were used for intracellular staining: Anti-human TBET FITC (1:50, clone 4B10, BioLegend, cat # 644812), Anti-human Granzyme B FITC (1:100, clone QA16A02, BioLegend, cat # 372206), Anti-human Granzyme K PE (1:25, clone GM26E7, BioLegend, cat # 370512), Anti-human FOXP3 PE CF594 (1:25, clone 236A/E7, BD Biosciences, cat # 563955), Anti-human GATA3 PE CF594 (1:25, clone L50-823, BD Bioscience, cat # 563510), Anti-human Amphiregulin PE Cy 7 (1:25, clone AREG559, Invitrogen, cat # 25-5370-42), Anti-mouse Nur77 APC (1:25, clone REA704, Miltenyi, cat # 130-111-231), Anti-human EOMES APC eF780 (1:25, clone WD1928, eBioscience, cat #47-4877-42), Anti-human RORgT BV650 (1:50, clone Q21-559, BD Biosciences, cat # 563424), Anti-human Perforin eF450 (1:100, clone dG9, Invitrogen, cat # 48-9994-42). Cells were analyzed on a BD LSR Fortessa flow cytometer. Data were plotted using FlowJo v10.7.1 (TreeStar) and Prism (Graphpad, v9).

#### PT cell culture

Commercially available human primary PTs from 6 donors (3 males and 3 females, Lonza Walkersville Inc) were expanded at passage 4, and studied at passage 5. The main donor characteristics are summarized in Supplementary Table 2. Cells were grown in custom-made Dulbecco’s modified Eagle’s medium (DMEM) containing 5.55mM D-glucose, 4mM L-glutamine, and 1mM sodium pyruvate, and supplemented with 10ng/mL human EGF, 0.05M hydrocortisone, 1x of Transferrin/Insulin/Selenium (Invitrogen), 10% v/v dialyzed fetal bovine serum (FBS), 50g/mL streptomycin, and 50units/mL penicillin, as previously^108, 109^. Cells were serum-starved for 24h prior to collection for gene expression, metabolite measurements, and assessment of metabolic function. For gene expression experiments, cells were washed with PBS, harvested with trypsin, and snap-frozen at -80°C until further analysis.

#### Assessment of metabolic function in human primary PT cells

Mitochondrial respiration was assessed in male and female PTECs by measuring their oxygen consumption rate (OCR) in a Seahorse XFe96 analyzer (Agilent). Glycolysis was also assessed by monitoring the extracellular acidification rate (ECAR). Upon 80-90% confluence, cells were detached with 0.25% trypsin (5min, 37°C), counted and seeded in a Seahorse XFe96 Cell Culture Microplate at a density of 15,000 cells/well in 100µL of DMEM complete media. After adhering for 6h, PT cells were exposed to serum starvation conditions for 24h. One hour prior to the metabolic function assay, cells were washed with phenol-free basal media (Agilent) and exposed to 150µL of assay media, which included 2mM glutamine and 5.55mM glucose. During the assay, OCR and ECAR were recorded at baseline and after metabolic stress. To induce metabolic stress, 25µL of oligomycin, p-trifluoromethoxy carbonyl cyanide phenyl hydrazone (FCCP), 2-deoxyglucose (2-DG), and Rotenone + Antimycin A (Rot+AA) were sequentially injected into the microplate wells. After optimization, the following working concentrations were stablished for each drug: oligomycin: 1µM; FCCP: 0.3µM, 2-DG: 100mM; Rot: 1µM; AA: 1µM. Basal respiration, ATP-linked respiration, maximal respiratory capacity, and reserve capacity were assessed by calculating the area under the curve (AUC) from OCR curves (**Fig. 3b, c**). Basal glycolysis, maximal glycolytic capacity, and glycolytic reserve were determined by calculating the AUC from ECAR curves (**Supplementary Fig. 6**).

#### Cell metabolite measurements

##### Sample preparation

Male and female primary PTs were grown on 6-well plates and subjected to starvation as described above. The levels of intracellular metabolites were then determined using liquid chromatography-mass spectrometry. After collecting the supernatant, 1mL of extraction solvent (80:20 mixture of methanol:water) was added into each well, in order to extract intracellular metabolites. Plates were placed on dry ice. The adherent material was then triturated, collected into Eppendorf tubes, and stored at -80°C. Cell lysate collection was followed by 3 freeze-thawing cycles in dry ice (to shift sample temperature between -80°C and -20°C). The insoluble material from each sample was then precipitated by centrifugation at full speed for 5min. The resulting pellet was dried at room temperature and used for total RNA quantification using the Quant-iT Ribogreen assay (Invitrogen). In turn, the metabolite extract was dried under high purity nitrogen gas (turbovap) and resuspended with appropriate volume of buffer (0.5µL of LC-MS grade water to 1µg of RNA) based on total RNA levels. The appropriate volumes of heavy-labelled (^13^C/^15^N) reference metabolites were spiked into each reconstituted sample for quantitation. The heavy-labelled metabolites used as internal reference standards were acquired in as a metabolite extract from yeast that had been 99% labelled with ^13^C-glucose and ^15^N-ammonia. To determine background metabolite signals, a mock plate without cells and equal volume of media was processed in parallel to the study plates.

##### Liquid chromatography-mass spectrometry (LC-MS)

Cellular metabolites were measured by injecting 2µL of sample in full scan MS1 mode using an Agilent 6550 qToF mass spectrometer coupled to an Agilent 1290 binary pump UPLC system. Most polar metabolite analytes presented here were measured using an Agilent ZORBAX ExtendC18 1.8 µm, 2.1 mm X 150 mm reverse phase chromatography using tributylamine as an ion paring agent as previously described^110^. The Agilent 6550 qToF was fitted with a dual AJS ESI source and an iFunnel with a gas temperature set to 150°C at 14L/min and 45psig. Sheath gas temperature was set to 325°C at 12L/min. Capillary and nozzle voltages were set to 2000V. Funnel conditions were changed from default to -30V DC, high pressure funnel drop -100V and RF voltage of 110V, low pressure funnel drop -50V and RF voltage of 60V. Metabolite annotation in full scan data was achieved by matching exact mass and retention time to an in-house database. The retention time and exact mass database were prepared by analyzing a collection of neat standards using the chromatographic method described above and confirming retention times by MS/MS fragmentation of neat standards.

##### Metabolite data analysis

Metabolite raw data was extracted directly from .d folders and integrated in profile mode using an R-based software package developed by the Rosebrock Lab; ChromXtractorPro (personal correspondence K. Laverty and A. Rosebrock). The metabolites whose intensity in all the study samples fell at or below their intensity in the blank (consisting of resuspension buffer only) were excluded from further analyses. Next, the integrated light (L) intensity of each metabolite was normalized to the intensity of its internal heavy (H) standard. The L/H ratio minimized the potential stochastic variation in the signal produced by the instrument due to changes in humidity and/or temperature, enabling the relative quantitation and comparative analysis of each metabolite. The analysis enabled the detection of 158 intracellular metabolites^111^. Data corresponding to the intracellular levels of NAD, β-nicotinamide mononucleotide, ATP, GTP, ITP, UTP were interrogated.

#### Gene expression validation studies

RNA was extracted from the cell pellets of human primary male and female PT cells using the RNAeasy Mini Kit (Qiagen). After quantifying RNA concentration in a Nanodrop instrument (Thermo), 300-700ng of RNA were retrotranscribed to cDNA using the High-Capacity cDNA Reverse Transcription Kit (Applied Biosystems). Male and female PTs had been grown and serum-starved as above. In these cells, gene levels of *KDM5D*, *UTY*, *EIF1AY*, *EIF1AX*, *DDX3X*, *MT1F*, *MT1G*, and *MT1H* were measured by real-time quantitative PCR using a Power SYBR® Green PCR Master Mix reagent (Applied Biosystems) and normalized to RPL31. The fluorescent signal was measured in a LightCycler® 480 Instrument II (Roche). All primer sequences employed in this study are summarized in **Supplementary Table 10**.

#### Quantification and statistical analysis

Statistical tests were conducted within R and using GraphPad Prism 9 software. For all comparisons, normality was determined using a Shapiro-Wilk test. Group-to-group differences were assessed using two-tailed unpaired T tests for variables following a normal distribution, and Mann-Whitney tests for variables with a non-parametric distribution. All p values below 0.05 were considered significant. Significance level for each test is indicated in the figures. For each experiment, n is reported in the figure legends and represents the number of samples.

#### Data availability

Count matrices from our complete data object are being submitted to NCBI GEO, and will be made publicly available upon publication. Additional information and data are available from the authors upon reasonable request, and in line with University Health Network (UHN) and UHN Research Ethics Board policies.

#### Code availability

We are preparing a Github repository for the custom scripts generated for data analysis.

## Supplementary Tables

**Supplementary Table 1.** ST1-Results of sex analyses. Results of genes identified with Varimax rotated PCA, sPLS-DA, and differential gene expression analysis using MAST comparing male and female proximal tubular cells.

**Supplementary Table 2.** ST2-Primary PT donor characteristics. Characteristics of the donors from which primary proximal tubular epithelial cells were isolated for metabolic studies.

**Supplementary Table 3.** ST3-GSEA significant results. Summary of significant gene set enrichment analysis terms between male and female proximal tubular cells.

**Supplementary Table 4.** ST4-DEGs LD NK & T cells Vs PBMC. Results of differential gene expression analysis using Seurat comparing kidney NK and T lymphocytes to circulating lymphocytes from two studies.

**Supplementary Table 5.** ST5-Cell cell interactions Omnipath. Results of aggregate cell cell communication inference with consensus and individual scores across methods, with Omnipath used as the reference interaction database

**Supplementary Table 6.** ST5-Cell cell interactions with complexes CellPhoneDB. Results of aggregate cell cell communication inference with consensus and individual scores across methods, with CellPhoneDB used as the reference interaction database

**Supplementary Table 7.** ST7-Patient characteristics. Characteristics of the study population.

**Supplementary Table 8.** ST8-CellRanger summaries of sequenced samples. CellRanger summaries with sample metrics for each sequenced sample.

**Supplementary Table 9.** ST9-Curated cell annotation file. Curated marker gene list for cell type annotations.

**Supplementary Table 10.** ST10-qPCR sequences. Primer sequences used for qPCR validation of sex differences in PT cells.

---

**Supplementary Figure 1:**
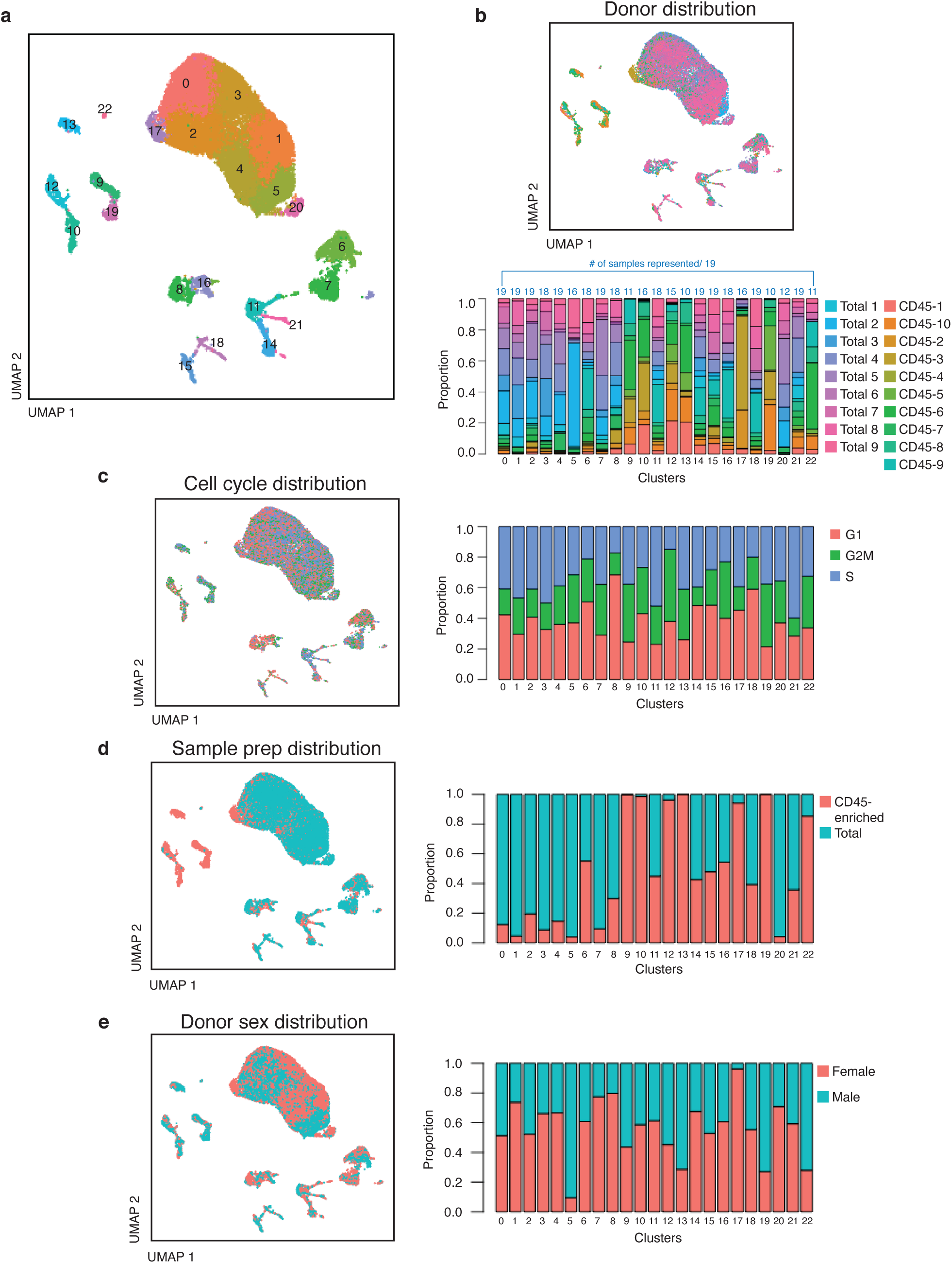
Additional proportion plots of total kidney dataset. (**a**) Clustering of total combined dataset of 27677 cells results in 23 clusters (**b**) Individual sample contribution to clustering, demonstrating that clusters are comprised of cells captured from multiple donors and in most cases all 19 samples contribute to each cluster. (**c**) Cell cycle assignment of clusters, with no exceptional variability in cell cycle state across clusters. (**d**) Distribution of sample preparation method (total homogenate versus CD45-positive magnetic bead enrichment) across clusters. **(e**) Distribution of donor sex across clusters.

**Supplementary Figure 2.**
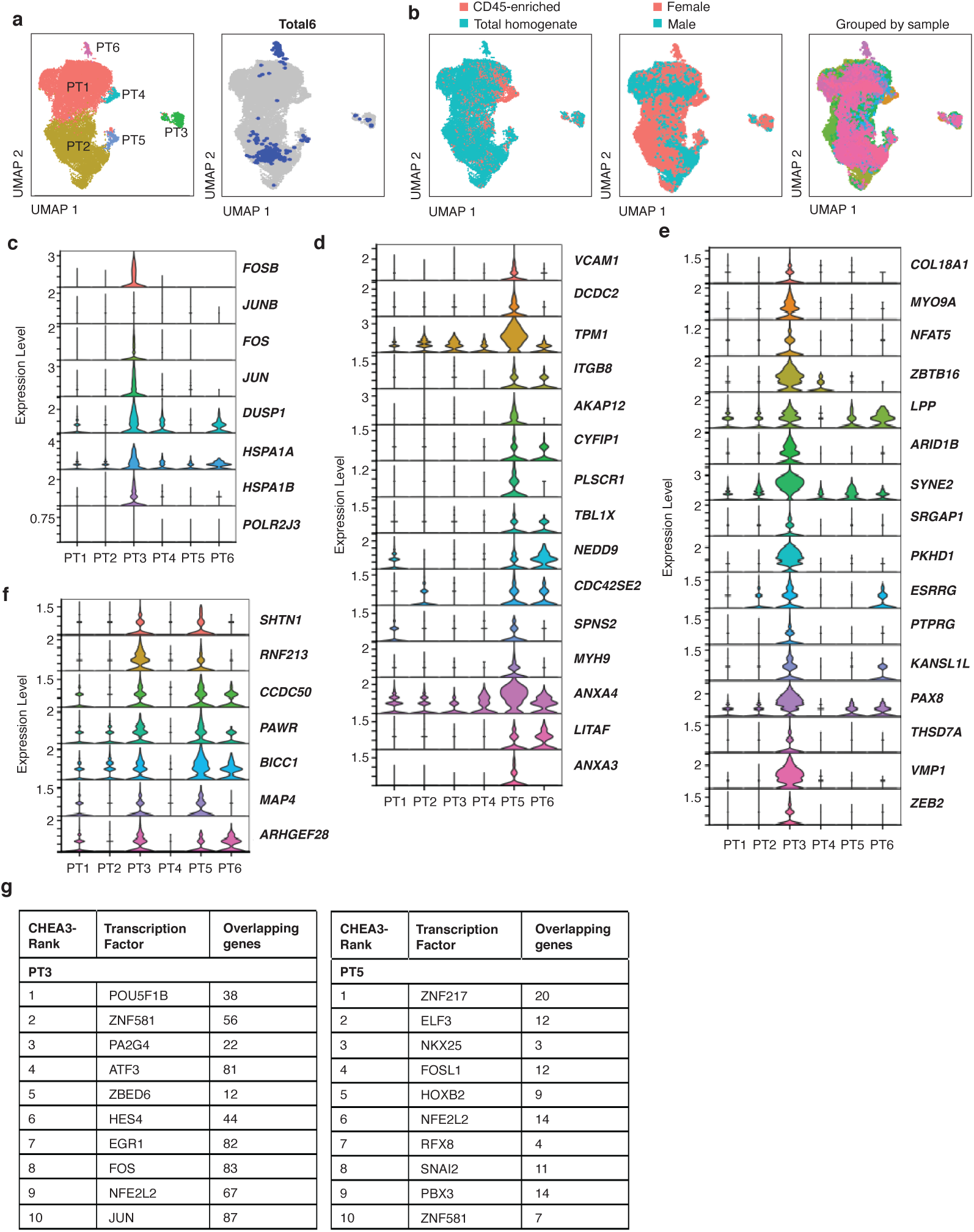
Heterogeneity within Proximal Tubular (PT) dataset. (**a**) Subclustering of PT dataset yielded 6 clusters; PT6 is predominantly composed of cells from one donor: “Total6”. (**b**) Distribution of sample preparation method, sex, and donor identity across the PT dataset; PT4 is composed of cells from CD45-enriched samples. (**c**) Stacked violin plots showing enrichment of dissociation stress markers in PT3. (**d**-**f**) Stacked violin plots showing markers of the ‘scattered tubular cell’ and ‘failed PT repair’ population enriched in PT5(**d**), PT3(**e**), and both PT3 and PT5(**f**). (**g**) Transcription factor analysis using CHEA3, which illustrates the top 10 transcription factors predicted to regulate PT3, and separately, PT5 genes.

**Supplementary Figure 3.**
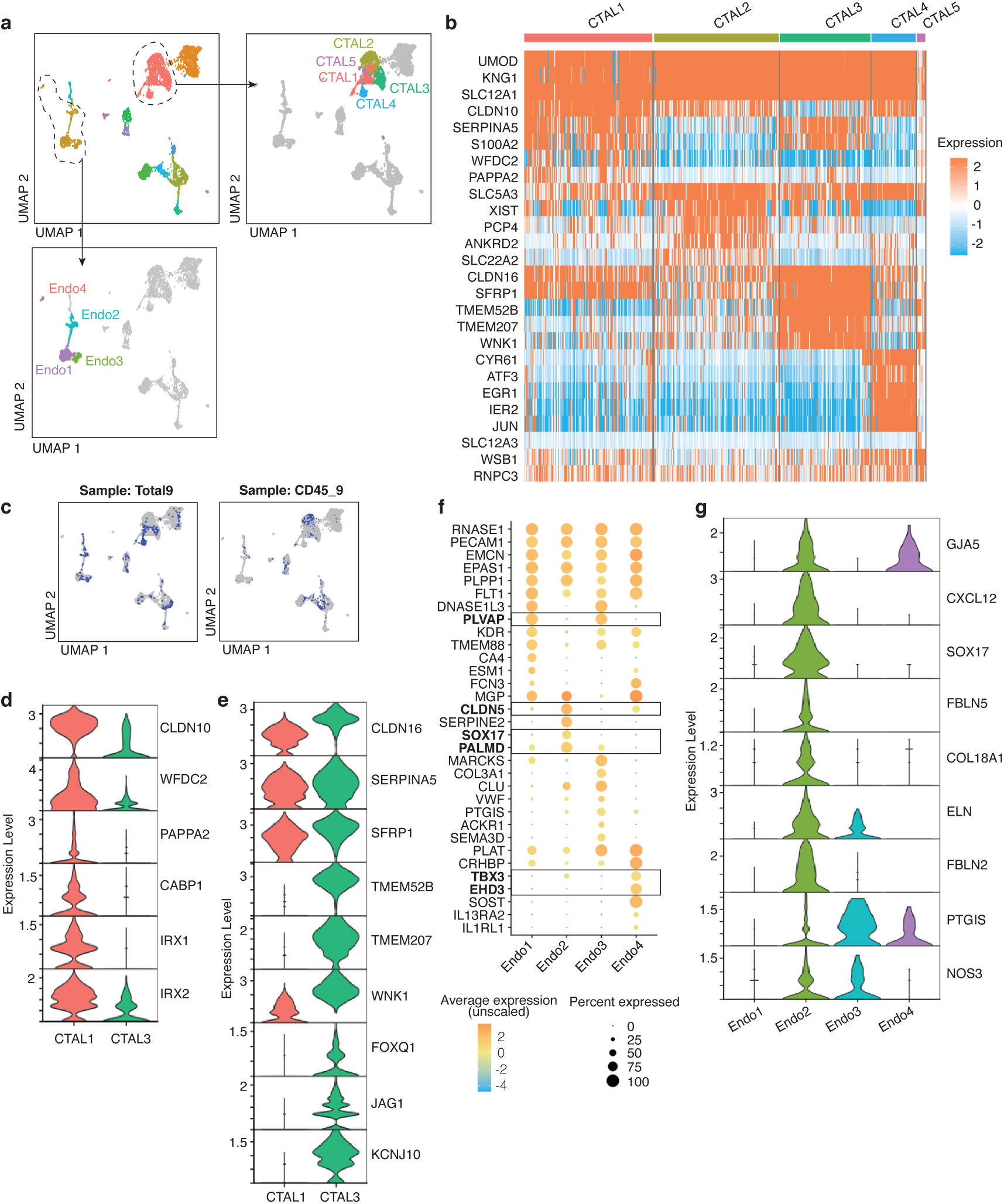
Heterogeneity in CTAL and Endothelial cell populations. (**a**) 5 CTAL clusters and 4 endothelial clusters were identified. (**b**) Heatmap depicting expression of the marker genes of CTAL1-5. (**c**) CTAL2 and 4 are each chiefly comprised of cells from one donor (Total9 and CD45_9, respectively). Selected marker genes of *CLDN10*-enriched CTAL1 (**d**) and *CLDN16*-enriched CTAL3 (**e**) populations, respectively. (**f**) Bubble plot showing enrichment for specific endothelial cell markers in all subpopulations; expression of peritubular capillary markers (*PLVAP, TMEM88, DNASE1L3*) in Endo1 and Endo3 respectively; expression of afferent arteriole and vasa recta genes (S*OX17*, *SERPINE2*, *CLDN5*, *CXCL12* and reduced *KDR*) in Endo2; and expression of glomerular microvascular endothelial cell markers in Endo4 (*EDH3*, *SOST* and *TBX3*).(**g**) Increased expression of extracellular matrix genes seen in Endo2 (characterised as afferent arterioles and vasa recta). Of the two peritubular populations described (Endo1 and Endo3), Endo3 is shown to have higher expression of vasodilators (*PTGIS* and *NOS3*) than Endo1. Endo4 illustrates expression of *GJA5* and *PTGIS*.

**Supplementary Figure 4.**
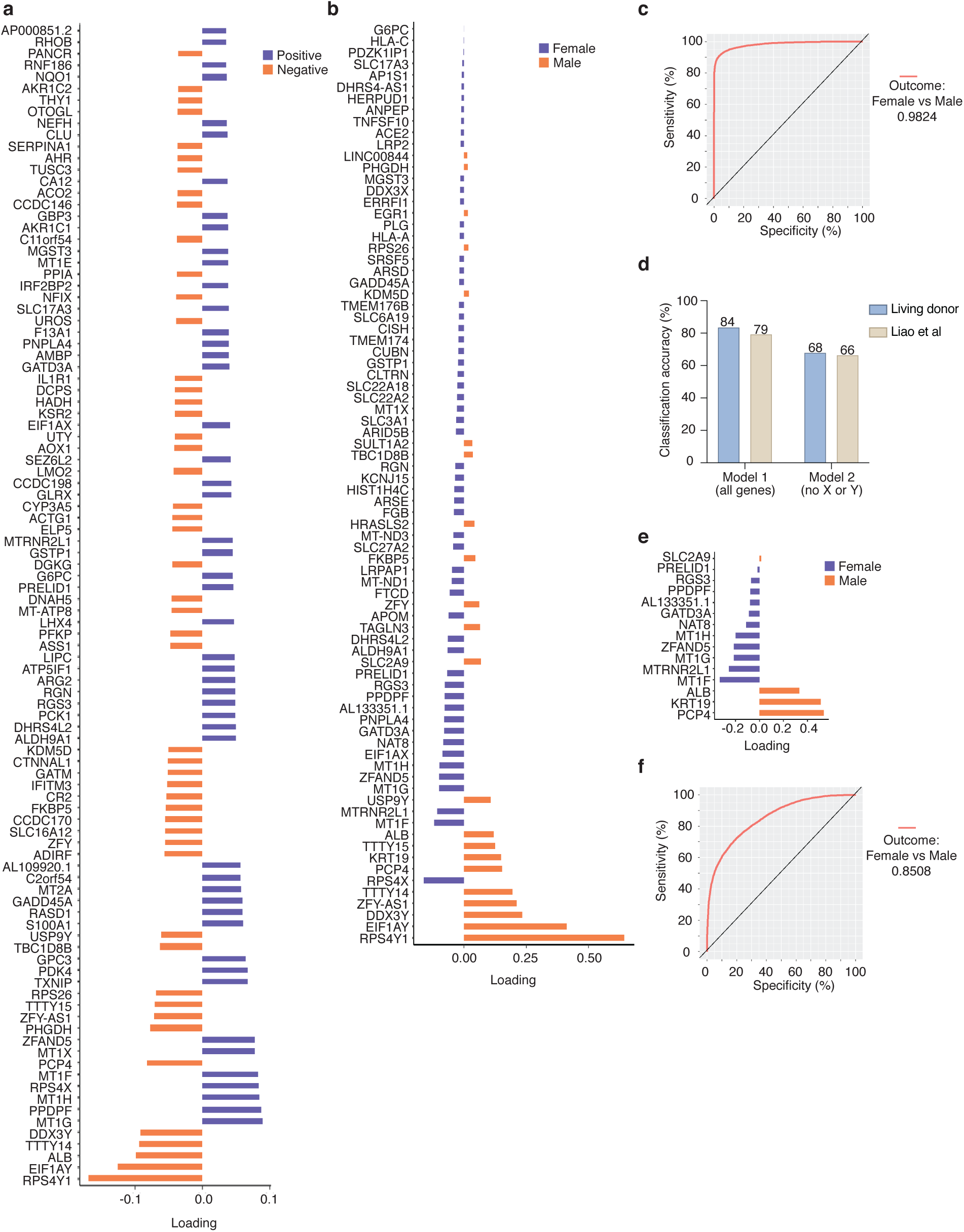
Varimax PCA and sparse partial least squares discriminant analysis (sPLS-DA) identifies sex differences in proximal tubular (PT) epithelial cells. (**a**) Top 100 genes (50 from each end of the component) associated with varimax-rotated principal component 12 which revealed sex differences in proximal tubule cells. (**b**) Plot of 80 genes that were selected as variables in the sPLS-DA classifier (Model 1) from all detected genes. (**c**) Receiver operating characteristic (ROC) curve from Model 1 predict male and female sex with accuracy of 98%. (**d**) Plot of 15 genes in Model 2 (using all detected genes except those encoded on X or Y chromosomes as input) where 15 genes were selected as variables in the classifier. (**e**) ROC curve from Model 2. (**f**) Barplot of classification accuracy using Model 1 versus Model 2 to classify PT cells of the living donor data and of a validation dataset from Liao *et al*.^20^

**Supplementary Figure 5.**
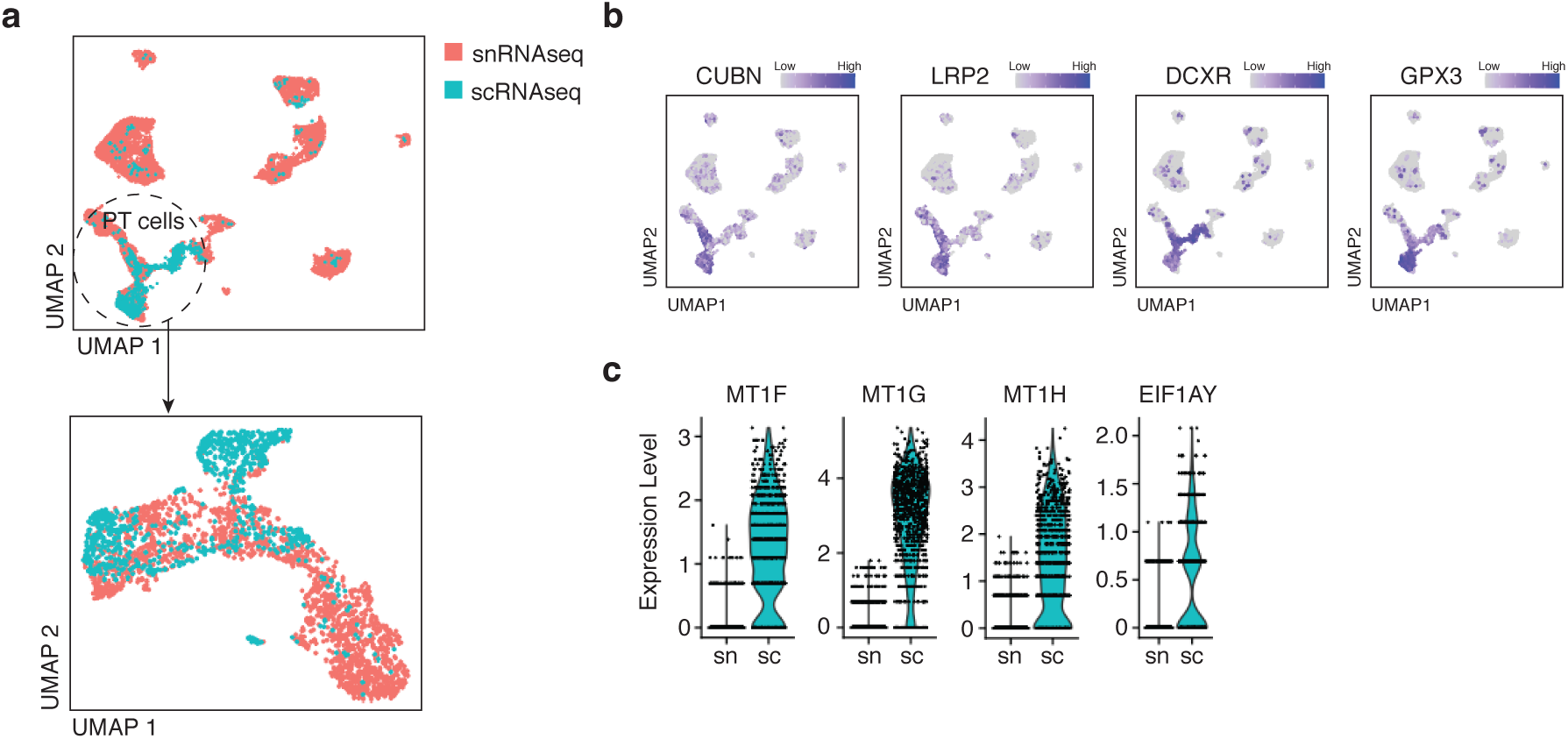
Comparison of single nucleus RNA sequencing and single cell RNA sequencing. (**a**) Data integration from a pilot sequencing experiment in which a single biopsy was divided and subjected to scRNAseq and single nucleus RNAseq (snRNAseq). From the integrated data, PT cell clusters were identified and analyzed. (**b**) Expression of PT cell marker genes used to identify clusters of PT cells in the integrated datasets. (**c**) Comparison of select genes from scRNAseq and snRNAseq reveals that several key genes exhibiting dichotomous expression across sexes as reported here are differentially captured by the two sequencing techniques.

**Supplementary Figure 6.**
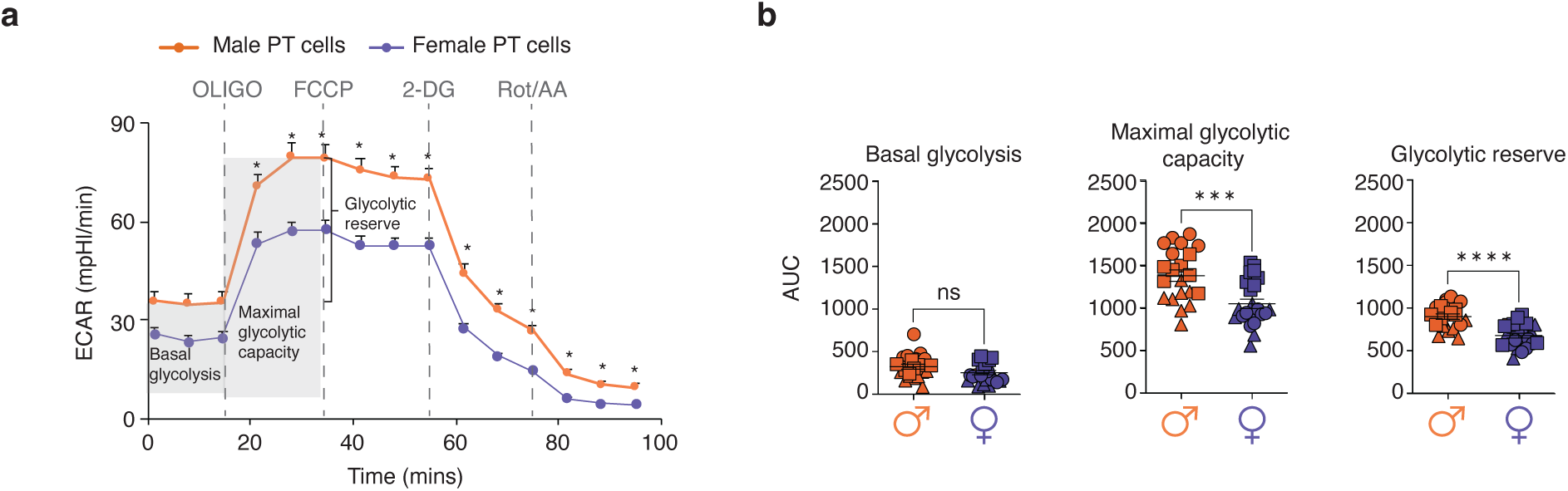
Sex differences in the glycolytic rate of proximal tubular (PT) cells. (**a**) The extracellular acidification rate (ECAR) was monitored to assess the glycolytic metabolism of male and female PT cells at baseline and after metabolic stress. To induce metabolic stress, the following sequence of drugs was injected: 1μM oligomycin, 0.3μM FCCP, 100mM 2-DG, 1mM Rot/AA. (**b**) The basal glycolysis (p=0.063, u=162), maximal glycolytic capacity (p=0.0004, t=3.832, df=42), and glycolytic reserve (p<0.0001, t=5.331, df=42) of male and female PT cells were calculated from the ECAR curves in (**a**) (n=3 donors/sex; n=6-8 replicates/donor). Group-to-group differences were assessed using two-tailed unpaired t-test for variables following a normal distribution, and Mann-Whitney tests for variables with a non-parametric distribution. *p<0.05; ***p<0.001; ****p<0.0001. PT, proximal tubule; AUC, area under the curve; ECAR, extracellular acidification rate; FCCP, p-trifluoromethoxy carbonyl cyanide phenyl hydrazone; 2-DG, 2-deoxyglucose; Rot, rotenone; AA: antimycin A.

**Supplementary Figure 7.**
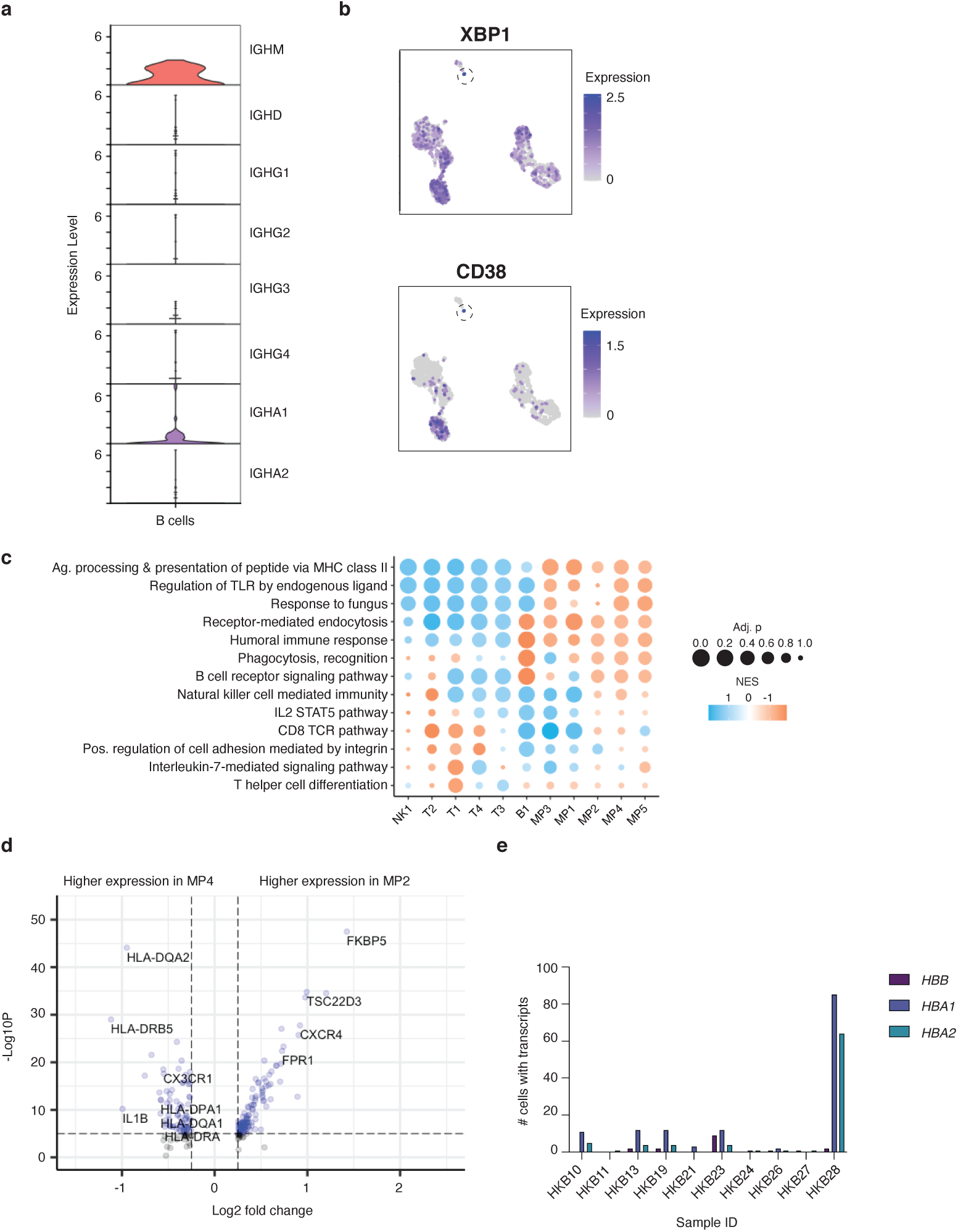
Additional immune cell phenotyping data. (**a**)Expression of immunoglobulin heavy chain genes within the B cell cluster, showing low abundance of class-switched B cells in living donor kidney. No IGHE transcripts were detected. (**b**) Very few plasma cells marked by high *XBP1* and *CD38* expression were identified. (**c**) Pathway analysis summary for immune populations, indicating an enrichment in cell-type specific pathways in support of cluster annotations. (**d**) Differential gene expression between two clusters (MP2 and MP4) of CD16^+^ monocyte-like cells identified an enrichment in antigen presentation genes in MP4, and differential expression of *CX3CR1* versus *CXCR4*. (**e**) Expression of hemoglobin transcripts in the CD45-enriched sequencing datasets, prior to any quality control thresholds or data cleanup steps. Sample HKB28 had the highest abundance of cells positive for hemoglobin transcripts, suggesting more circulating cells in this sample.

**Supplementary Figure 8.**
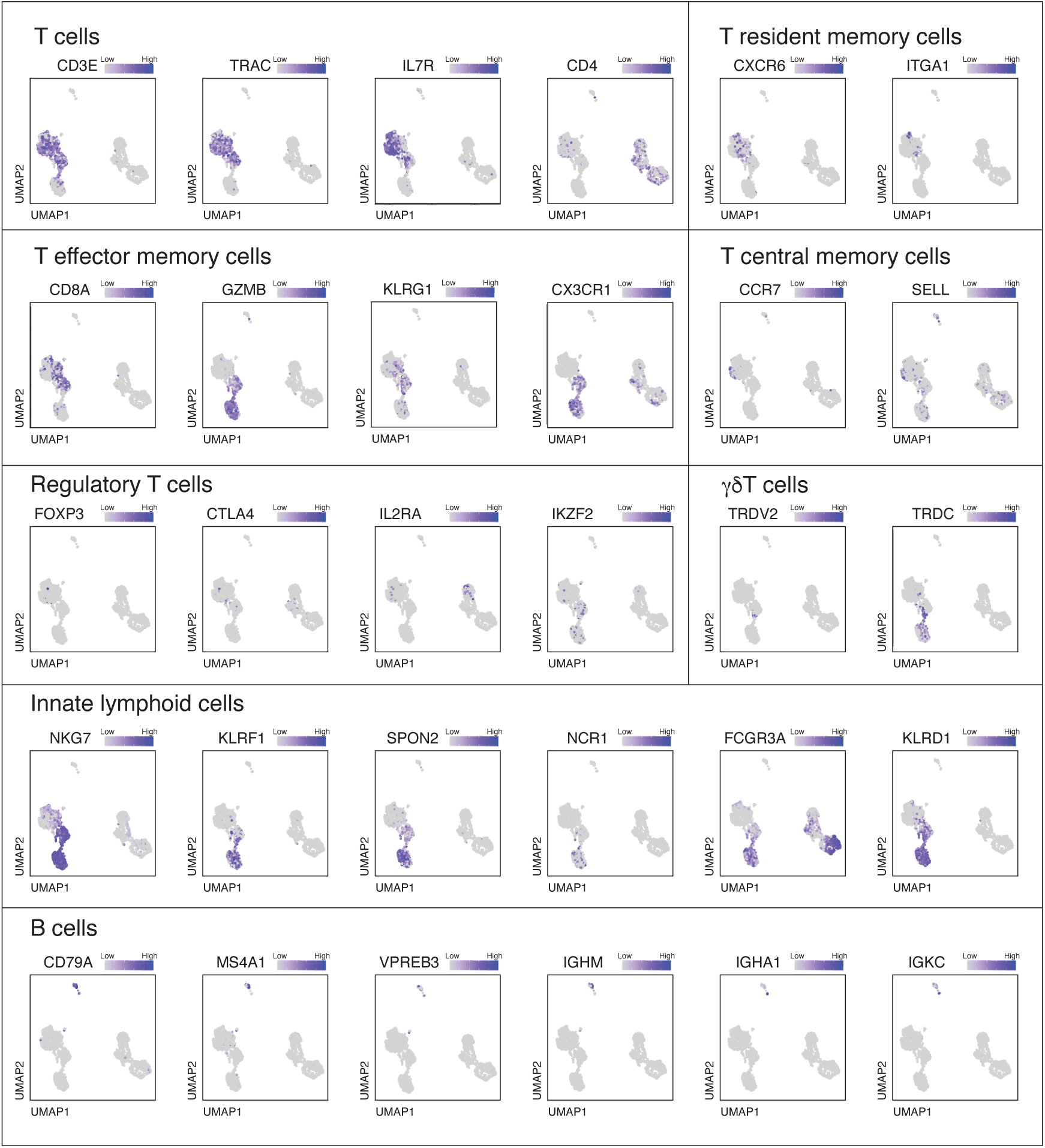
Annotation of lymphocyte populations. Additional feature plots used to annotate of lymphocyte cell types including general T cell markers and subset-specific markers of T resident memory, T effector memory, and T central memory cells, as well as markers of regulatory T cells, γδT cells, innate lymphoid cells and B cells.

**Supplementary Figure 9.**
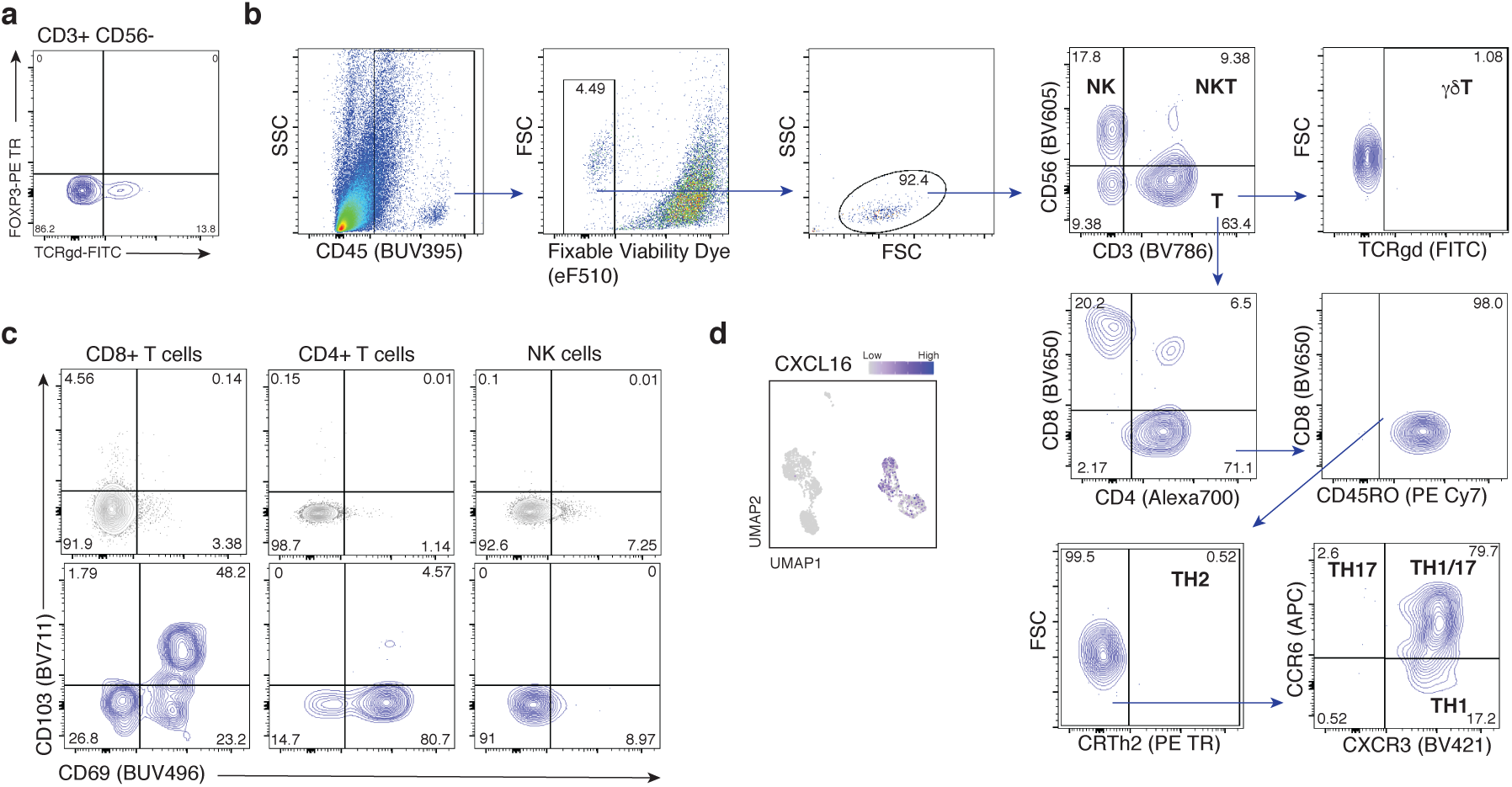
Additional supporting data for the identification of resident signatures in kidney lymphocytes. (**a**) No FOXP3 expression was noted on T cells, and TCRγδ staining validated the presence of γδT cells within healthy kidney. (**b**) Gating strategy for the identification of T helper subsets. (**c**) Co-expression of CD69 and CD103, characteristic of Trm cells on CD8^+^ and CD4^+^ T cells and NK cells of the blood (grey, top row) versus kidney (blue, bottom row). (**d**) Expression of the chemokine CXCL16 in myeloid cells of the kidney supporting recruitment of CXCR6^+^ lymphocytes.

**Supplementary Figure 10.**
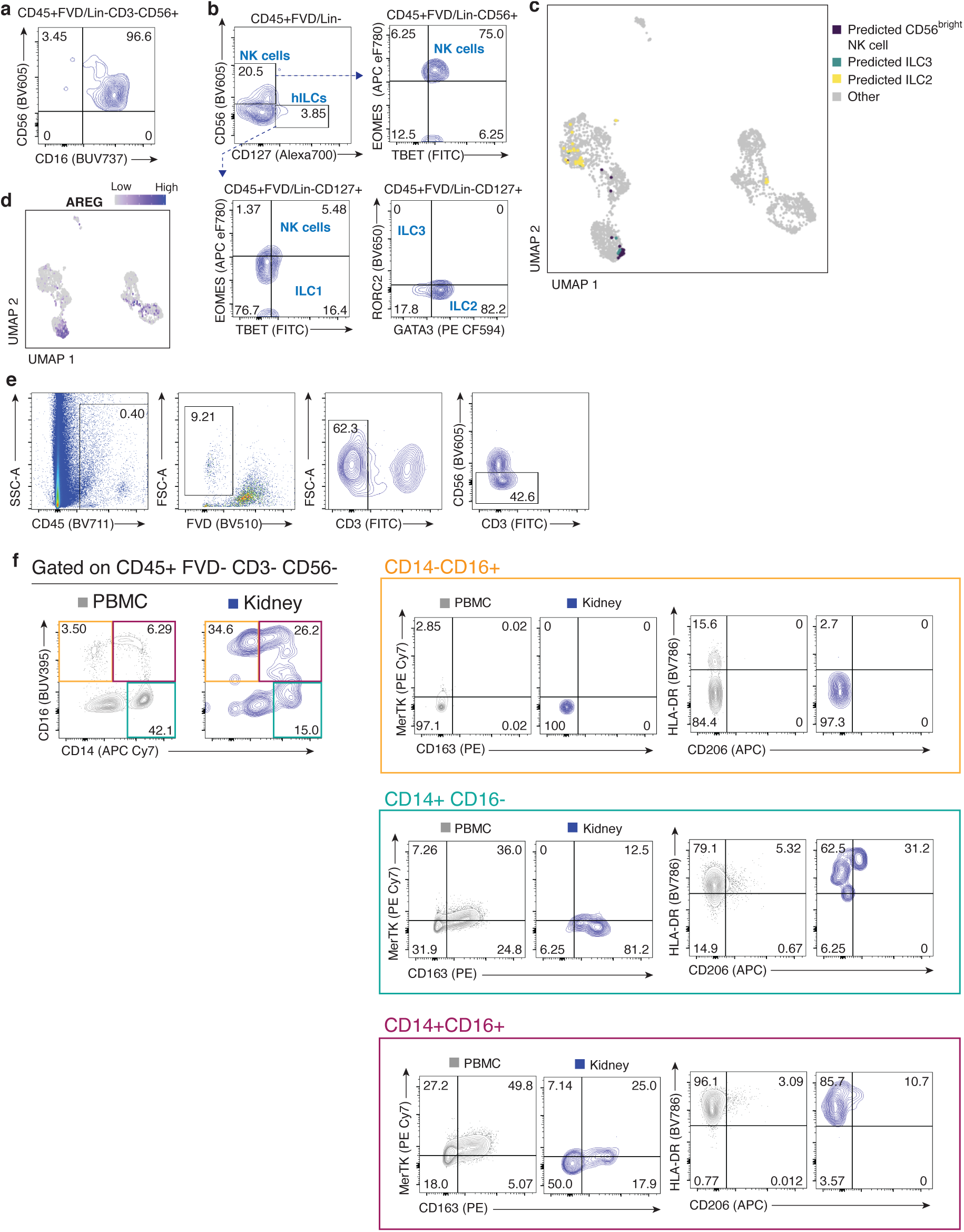
Identification of innate lymphoid cell and myeloid populations in healthy human kidney. (**a**) The majority of NK cells within kidney are CD56^dim^CD16^+^, while (**b**) helper ILCs are present in very low abundance in kidney tissue. (**c**) Predictive identification of CD56^bright^CD16^-^ NK cells, ILC3s, and ILC2s within kidney immune transcriptomic data. (**d**) High expression of *AREG* encoding amphiregulin in kidney NK cells. (**f**) Gating strategy to remove lymphocytes from the population of interest. (**f**) Relative to blood, kidney tissue is enriched in CD16^+^ myeloid populations, and also allowed for identification of a CD14^+^ CD206^+^HLA-DR^+^ population likely representing MP1.

**Supplementary Figure 11.**
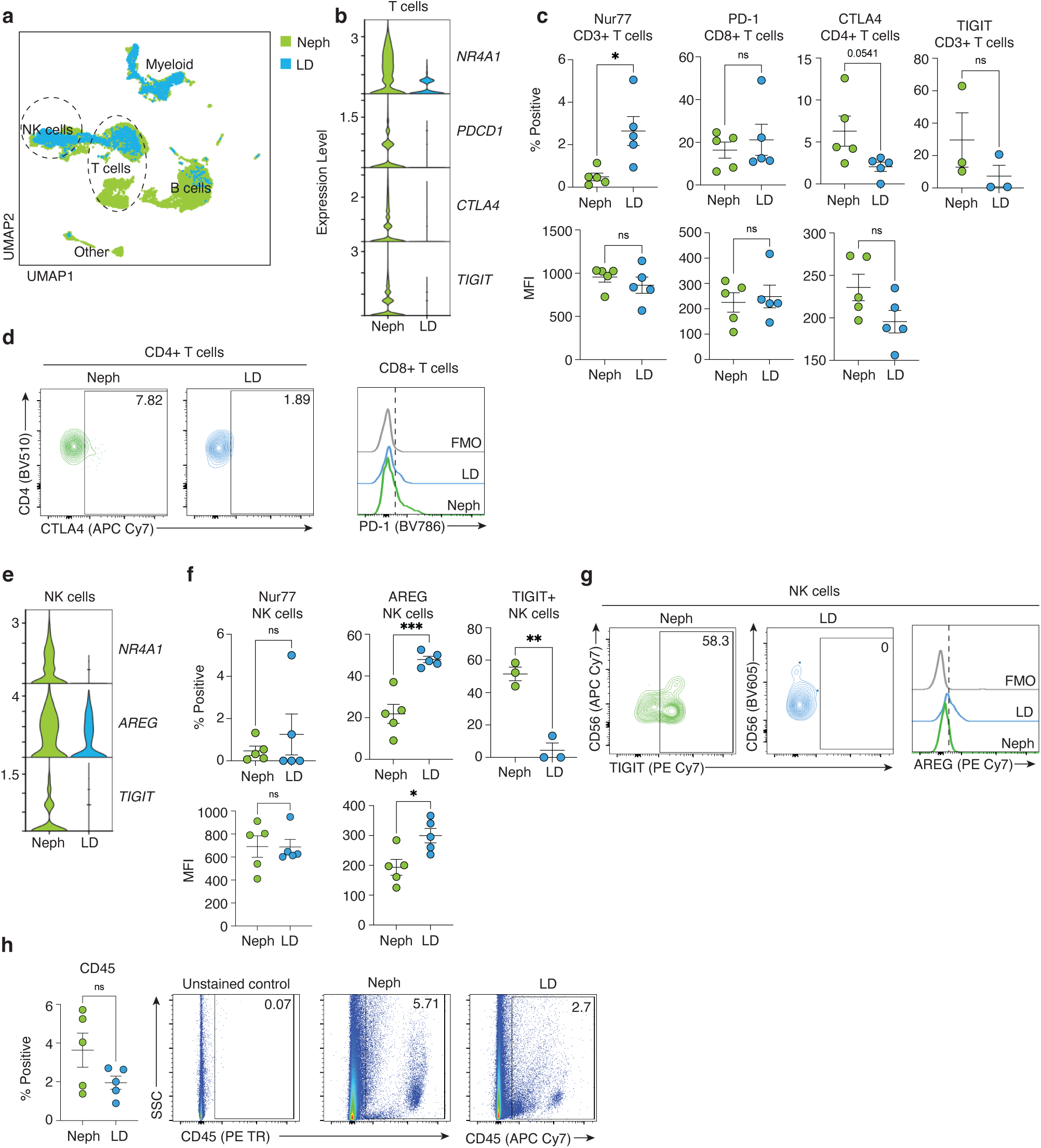
Comparison of sequencing data from nephrectomy versus living donor kidney specimens. (**a**) Integrated UMAP of kidney immune cells highlighting the contributions of cells derived from living donor versus nephrectomy tissue. (**b**) Within the T cell compartment, the activation marker *NR4A1* (encoding Nur77) along with checkpoint molecules *PDCD1 (*encoding PD-1), *CTLA4* and *TIGIT* were more highly expressed in nephrectomy data. (**c**), NR4A1 percent positivity (p=0.0152, t=3.076, df=8) and MFI (p=0.4206, u=8) on CD3^+^ T cells, PD-1 percent positivity (p=0.6905, u=10) and MFI (p=0.7024 t=0.3961 df=8) on CD8^+^ T cells, CTLA-4 percent positivity (p=0.0541, t=2.256, df=8) and MFI (p=0.0851, t=1.964, df=8) on CD4^+^ T cells and TIGIT percent positivity (p=0.2833, t=1.238, df=4) on CD3+ T cells were compared between living donor and nephrectomy-derived T cells. (**d**) Representative plots of CTLA-4 on CD4+ T cells and PD-1 on CD8+ T cells of living donor and nephrectomy-derived cells. (**e**) NK cells exhibited similar trends at the transcript level with higher *NR4A1*, *AREG*, and *TIGIT* gene expression in nephrectomy data. (**f**) While Nur77 protein was not differentially detected by percent positivity (p=0.5397, u=9) or MFI (p>0.999, u=12), AREG was higher in living donor NK cells by percentage (p=0.0006, t=5.420, df=8) and MFI (p=0.0182, t=2.959, df=8), and TIGIT (p=0.0015, t=7.728, df=4) was more highly detected on nephrectomy NK cells as shown in representative plots in **g**. (**h**) CD45+ cell elevation in nephrectomy samples did not reach significance (p=0.1129, t=1.780, df=8), however, increased immune cell (CD45^+^) abundance was observed in 3/5 nephrectomy samples tested, with high donor heterogeneity in immune cell abundance was observed, indicative of greater differences in tissue microenvironment between nephrectomy specimens. Group-to-group differences were assessed using two-tailed unpaired t-test for variables following a normal distribution, and Mann-Whitney tests for variables with a non-parametric distribution. *p<0.05; **p<0.01;***p<0.001; ****p<0.0001. Neph= nephrectomy, MFI=Median Fluoresence Intensity.

**Supplementary Figure 12.**
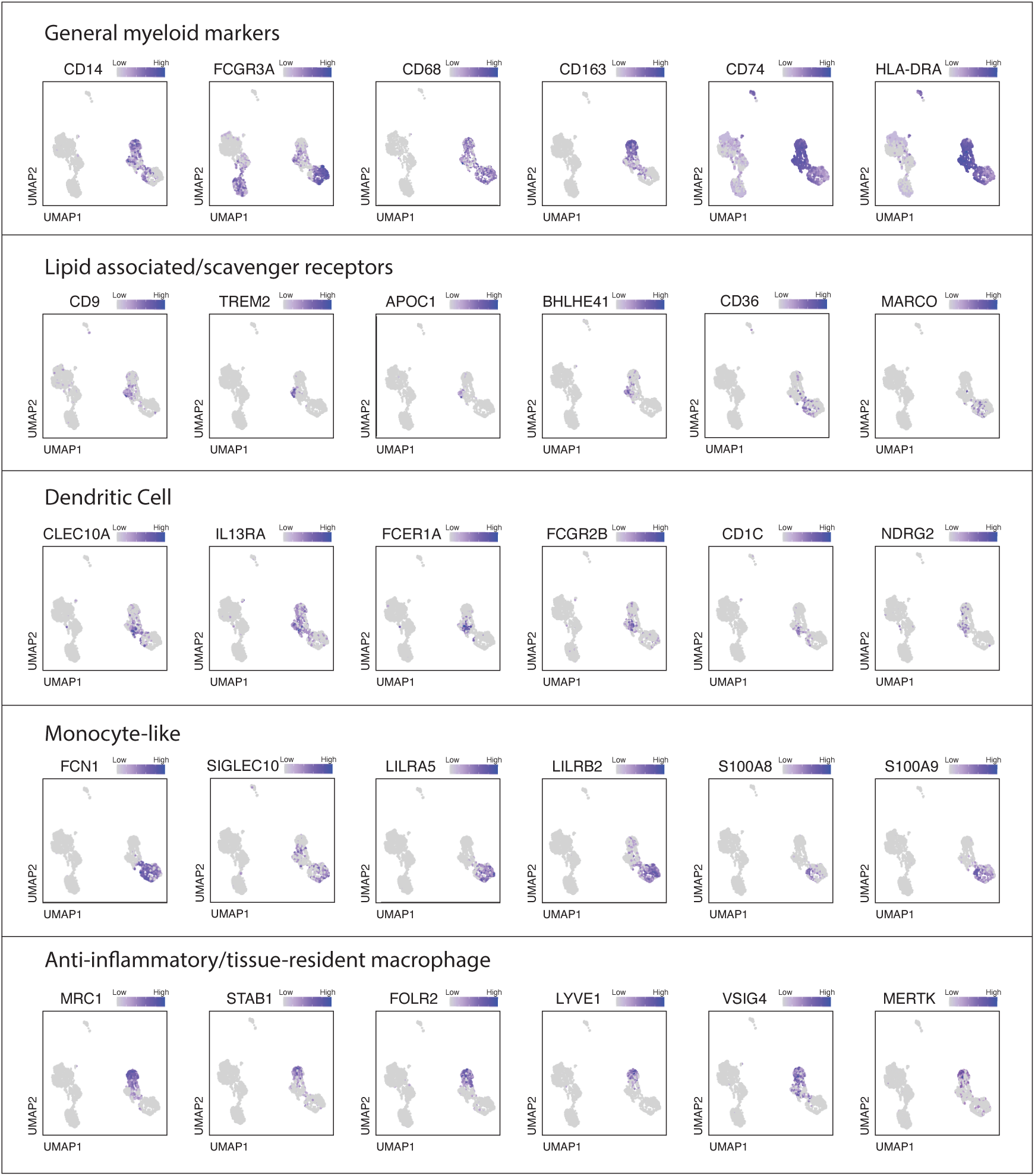
Annotation of myeloid populations. Additional feature plots of myeloid cells supporting cell type annotations, highlighting general myeloid lineage markers, expression of scavenger receptors, and markers of dendritic cells, monocytes and macrophages.

**Supplementary Figure 13.**
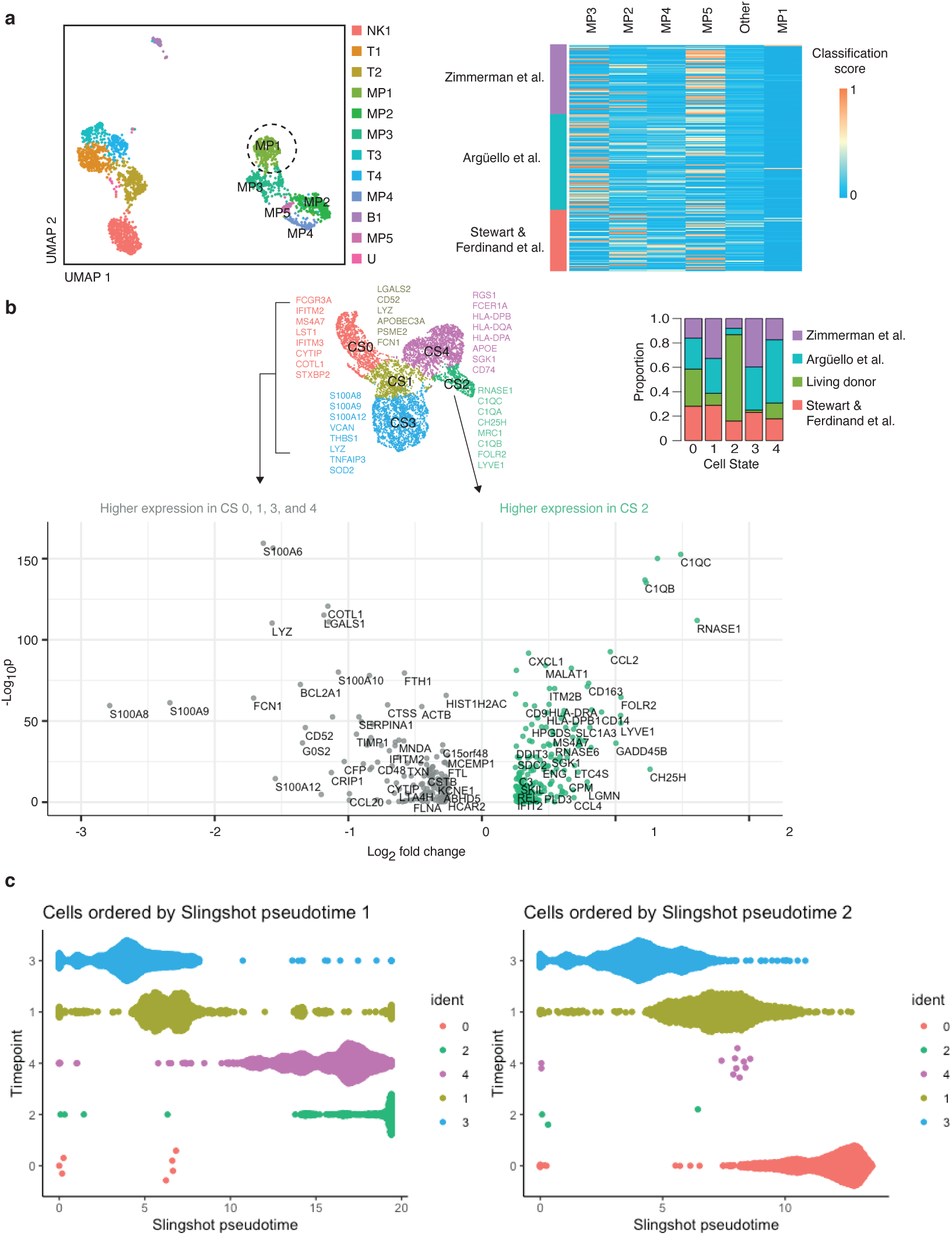
Additional supporting plots for macrophage cell state identification. (**a**) Based on our identification of 5 clusters of myeloid lineage cells in living donor kidney, we used SingleCellNet to classify cells from previously published datasets^3, 45, 46^ into our 5 cluster framework. Most cells captured in prior studies were classified as MP5 (CD14^+^ monocytes), the smallest cluster in living donors; while MP1 (circled) the largest cluster in living donor data was scarcely represented in previously published data. (**b**) Merging the three datasets specified in (**a**) with our living donor dataset confirmed 5 cell states (CS) where living donor data comprised the majority of CS2. A volcano plot depicts genes enriched in CS2 versus the remaining four cell states, supporting that CS2 represents a resident alternatively-activated tissue macrophage population that is uniquely enriched in living donor kidney tissue. (**c**)Slingshot pseudotime analysis supporting the annotation of CS1 as a transitional myeloid population across two suggested trajectories which placed CS2 and CS0 as the potential trajectory endpoints.

**Supplementary Figure 14.**
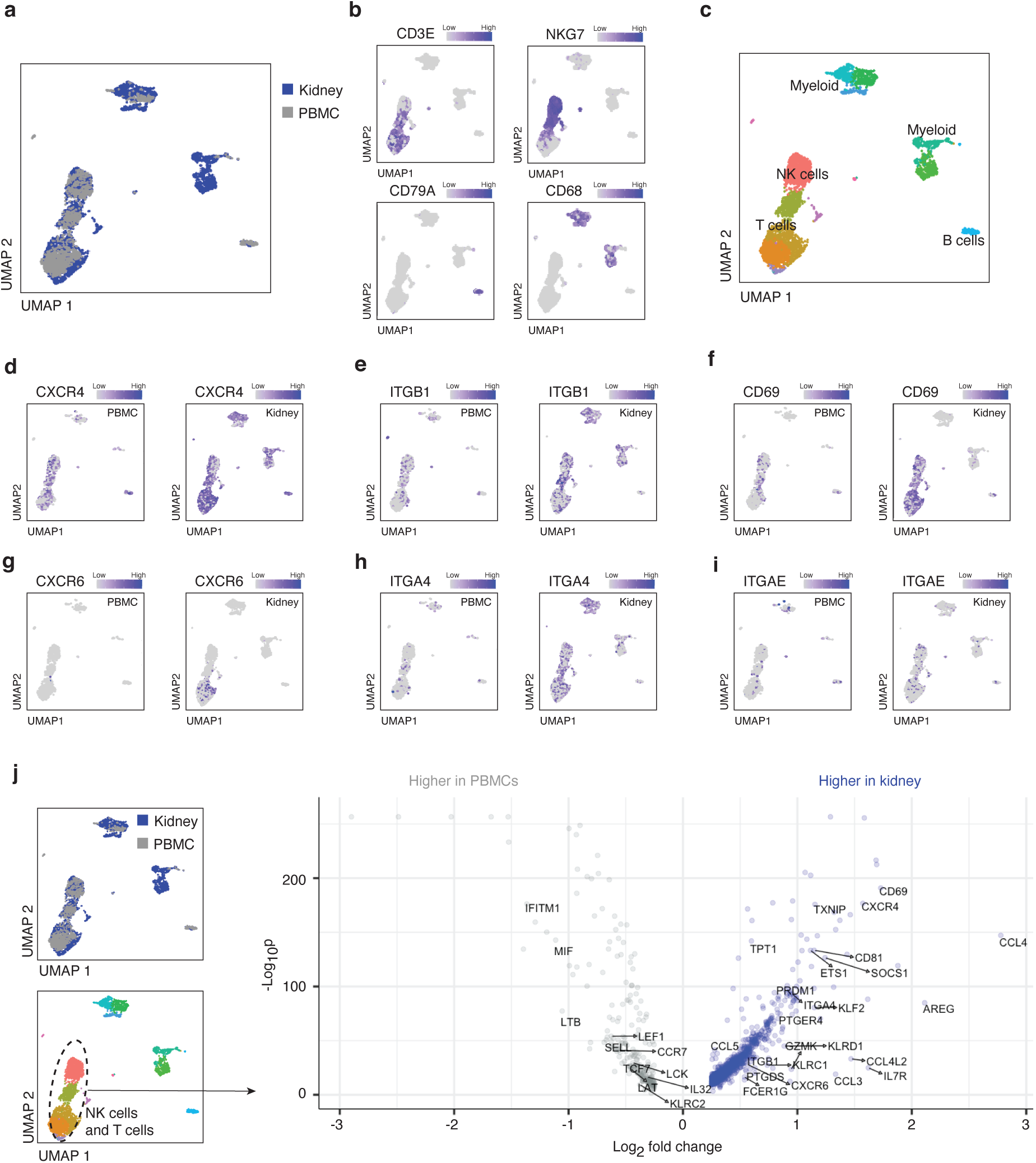
Integration of PBMCs and kidney immune single-cell data. (**a**) scRNAseq data from living donor kidney immune cells and PBMCs^48^ were integrated using Harmony. (**b**) Feature plots demonstrating expression of *CD3E*, *NKG7*, *CD79A*, and *CD68* used to annotate major immune populations in the combined dataset. (**c**) Annotation of major immune populations including T cells, NK cells, B cells and myeloid cells in the integrated PBMC and kidney immune dataset. Feature plots showing gene expression in PBMCs versus living donor kidney data of marker genes used for validation at the protein level including (**d**)*CXCR4*, (**e**)*ITGB1,* (**f**)*CD69,* (**g**)*CXCR6,* (**h**)*ITGA4* and (**i**)*ITGAE*. (**j**) Differential expression analysis of the T cells and NK cell clusters identifies genes which are upregulated in kidney lymphocytes and may represent kidney-adapted gene expression of these cells.

**Supplementary Figure 15.**
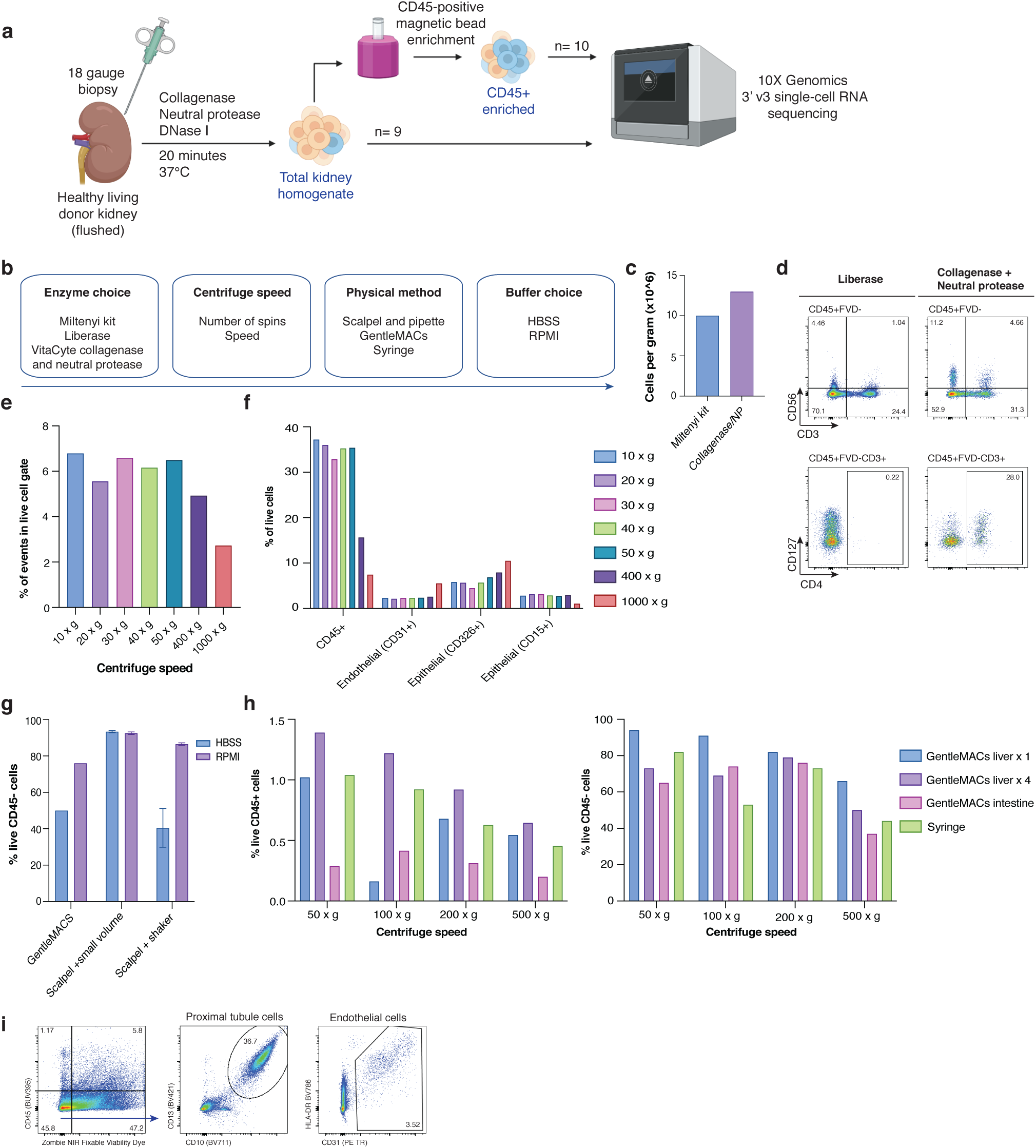
Optimization of kidney tissue digestion protocol. (**a**) Final experimental protocol for generating single-cell RNA sequencing data from living donor kidney. (**b**) Workflow of options tested in determining the optimal digestion method. (**c**) Using mouse tissue, a commercial Miltenyi kidney digestion kit was compared to a collagenase and neutral protease mixture to compare yield and viability, with collagenase and neutral protease demonstrating superior yield and comparable viability. (**d**) Using human nephrectomy tissue, Liberase was compared with collagenase and neutral protease, and flow cytometry was used to determine viability and cell phenotype, where it was determined that collagenase/neutral protease preserved key surface markers that appear to be cleaved by Liberase. (**e**) Fractions of dissociated human nephrectomy were centrifuged at different speeds to determine cell viability, which was reduced beyond speeds of 400 x g. (**f**) Using markers of key cell populations, by flow cytometry the contribution of different cell populations to each fraction by differential centrifugation determined that cell types were captured proportionally up to speeds of 400 x g. (**g**) To optimize yield and preservation of parenchymal cell viability, digestion in either HBSS or RPMI medium with collagenase and neutral protease was tested alongside physical methods of dissociation including using GentleMACs, a scalpel and a small volume of dissociation medium (n=2), and a scalpel with incubation with constant agitation in a shaker (n=2). Over all methods, RPMI preserved parenchymal viability better than HBSS, while overall the greatest viability was in using a scalpel and small volume of dissociation medium. (**h**) Viability of immune (CD45^+^) and parenchymal (CD45^-^) populations across physical methods and centrifuge speeds to test whether the relative abundance of cell population viability changes with more aggressive physical dissociation, where generally more gentle dissociation preserved parenchymal cell viability whereas more aggressive physical dissociation improved yield of immune cells. Different Gentlemacs™Tissue dissociator settings named based on organ optimized for were tested (liver, intestine, etc). No clear change in fractionation was observed in differential centrifugation of the samples. n=1 unless otherwise specified. (**i**) Flow cytometric analysis of kidney parenchymal cells, depicting the capture of live proximal tubular epithelial cells (CD10^+^CD13^+^) and endothelial cells (CD31^+^HLA^-^DR^+/-^).

**Supplementary Figure 16.**
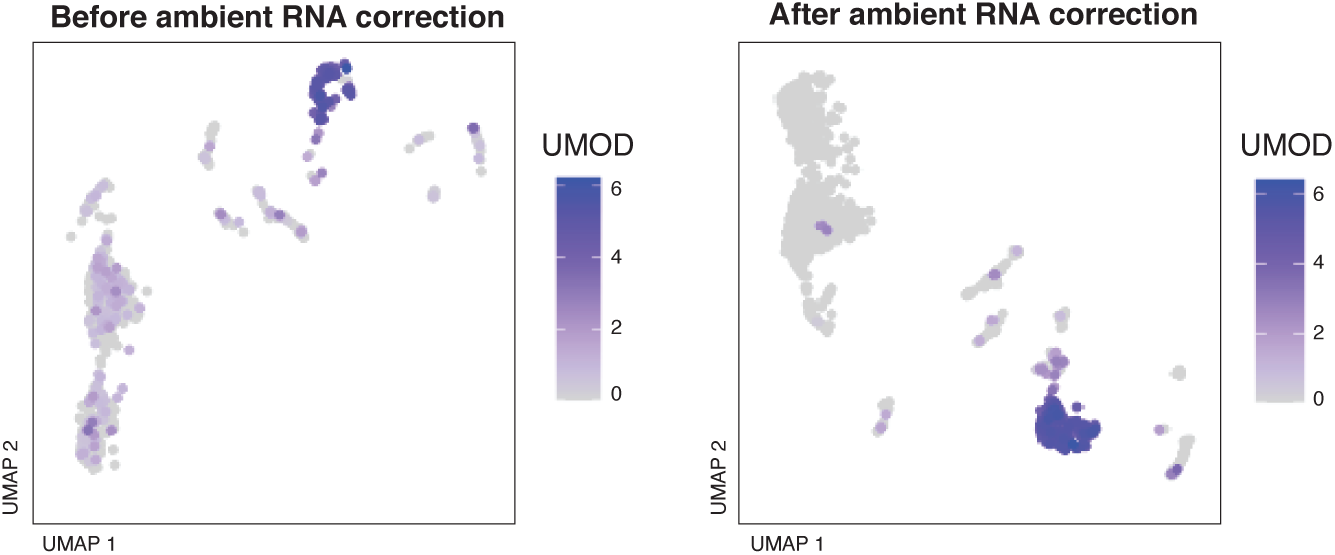
Ambient RNA contamination. Feature plots showing the expression of UMOD, a gene specific to cTAL/LOH cells, with widespread low level expression present across all clusters prior to ambient RNA correction and more biologically appropriate expression patterns after ambient RNA correction, demonstrated with a sample dataset (Total9).

## Notes

### Competing Interest Statement

The authors have declared no competing interest.

